# Integration of Transcriptome and Small RNA Sequencing to Decipher Molecular Interaction of *Chenopodium quinoa* Varieties with Cucumber Mosaic Virus

**DOI:** 10.1101/2020.07.20.212795

**Authors:** Nourolah Soltani, Margaret Staton, Kimberly D. Gwinn

**Affiliations:** Department of Entomology and Plant Pathology, University of Tennessee, Knoxville, TN, USA; Department of Plant Pathology, University of California-Davis, Davis, CA, USA

**Author notes:** Correspondence (KDG); (MS).

**Keywords:** *Chenopodium quinoa*, cucumber mosaic virus, RNASeq, small RNA, triterpenoid saponin, virus infection

## Abstract

Saponins are secondary metabolites with antiviral properties. Low saponin (sweet) varieties of quinoa (*Chenopodium quinoa*) have been developed because seeds high in saponins taste bitter. The aim of this study was to elucidate the role of saponin in resistance of quinoa to *Cucumber mosaic virus* (CMV). Differential gene expression was studied in time-series study of CMV infection. High-throughput transcriptome sequence data were obtained from 36 samples (3 varieties × +/- CMV × 1 or 4 days after inoculation × 3 replicates). Translation, lipid, nitrogen, amino acid metabolism, and mono- and sesquiterpenoid biosynthesis genes were upregulated in CMV infections. In ‘Red Head’, (bitter), CMV-induced systemic symptoms were concurrent with downregulation of a key saponin biosynthesis gene, TSARL1, four days after inoculation. In local lesion responses (sweet and semi-sweet), TSARL1 levels remained up-regulated. Known microRNAs (miRNA) (81) from 11 miR families and 876 predicted novel miRNAs were identified. Differentially expressed miRNA and short interfering RNA clusters (24nt) induced by CMV infection are predicted to target genomic and intergenic regions enriched in repetitive elements. This is the first report of integrated RNASeq and sRNASeq data in quinoa-virus interactions and provides comprehensive understanding of involved genes, non-coding regions, and biological pathways in virus resistance.

## 1. Introduction

Quinoa (*Chenopodium quinoa* Willd.) is a pseudocereal crop with high nutritional value [1, 2]. Quinoa varieties are classified as bitter or sweet depending upon saponin content of the seed. Although saponins in quinoa are highly accumulated in the seed pericarp, they are also present in lower concentrations in leaves. Saponin biosynthesis in quinoa is regulated by the triterpene saponin biosynthesis activating regulator-like1 (TSARL1) gene [3-5]. Mechanical abrasion or water rinsing is typically used to remove the saponin from the seed of bitter varieties [6, 7]. Because these methods are costly and reduce nutritional value of the seeds [6, 8], quinoa cultivars with low seed saponin content (sweet cultivars) are desired [9]. Saponins from quinoa are active against several plant pathogens including viruses and are pre-existing defense factors in other pathosystems [10-14]. Influence of saponin content on tolerance and/or susceptibility to viral diseases has not yet been investigated.

Plant viruses are obligate parasites that are dependent on their host to establish a successful infection cycle including gene expression, genome replication, protein synthesis, and intercellular movement. During the infection, protein-protein or protein-nucleic acid interactions may induce plant defense mechanisms such as innate immunity, translational repression, autophagy-mediated or ubiquitinated-mediated protein degradation, and RNA interference (RNAi) [15, 16]. Activation of the host immune responses results in either a compatible or incompatible response in the plant and is followed by a susceptible or resistance reaction to virus, respectively. Among these interactions, RNAi is one of the major evolutionarily-conserved defense mechanisms against viruses, and viral evasion from this resistance response is a crucial event in viral pathogenesis [17].

The RNAi mechanism induces gene silencing by targeting genes in a sequence-specific manner through chromatin modification, mRNA degradation or translation inhibition [18]. The RNAi mechanism is necessary to the host since it controls endogenous expression and translation and counteracts exogenous particles such as transposons and viruses [19, 20]. The mechanism of action in RNAi involves processing of double-stranded RNA (dsRNA) into 21-24 nucleotides by Dicer-like (DCL) enzymes [21]. Argonaute (AGO) family proteins cleave dsRNA into two single-stranded RNAs (guide and passenger RNAs), stabilize association of guide RNA onto the RNA-induced silencing complex (RISC), and direct the RISC toward the target sequence, which ultimately leads to transcriptional or posttranscriptional gene silencing [22]. Furthermore, gene silencing can be reinforced by incorporation of RNA-dependent RNA polymerase (RDR) enzyme activity through generation of secondary dsRNA that can result in systemic gene silencing [21, 23]. During viral infection in plants, recruitment of AGO1-5 and RDR1,6 plays an important role in viral gene silencing [24-26].

Small RNAs (sRNA) produced by plants are classified into micro (mi)RNA and short interfering (si)RNA that are cleaved to 21-24 nucleotideswith a preferential 5’ terminal base [24]. The miRNAs have incomplete complementarity with their targets and are cleaved through incorporation of DCL1 and AGO1 [24, 27, 28]. The siRNAs are excised from dsRNA that have perfect base-pairing with their target and are processed by association of different DCL, AGO, and RDR components [21]. Natural antisense siRNAs (nat-siRNAs) are generated through cleavage of endogenous dsRNA into 21 or 22nt by DCL4 or DCL2, respectively [29]. Heterochromatic siRNAs (hc-siRNAs) are comprised of 24nt dsRNA via cleavage of DCL3 and mediating of AGO 4/6/9. Hc-siRNAs are involved in silencing of heterochromatic, repetitive regions, coding sequences, gene promoter regions, and transposable elements via RNA-directed (PolIV) DNA methylation and histone modification. RDR2 is also recruited in the process to generate dsRNA to enhance siRNA biogenesis [24, 29-35]. Determination of differentially expressed miRNA and/or siRNA upon virus infection identifies major candidate small RNAs silencing viral genes. These candidates in combination with transcriptome profiles provide a comprehensive source of genes, miRNAs, and siRNAs as potential targets for enhancing plants resistance against viruses.

Transcriptome profile of *C. amaranticolori* (a weed host) and *C. quinoa* infected with different viruses determined modulation of photosynthesis, hormone signaling, plant-pathogen interaction, secondary metabolites, lipid, amino acid, protein, and carbohydrate metabolism [36-39]. However, there is no knowledge on the impact of virus infection of quinoa varieties differing in saponin levels on transcriptome and small RNA profiles of quinoa. Therefore, in this study, quinoa leaves from varieties with different seed saponin content were inoculated with cucumber mosaic virus (CMV), and differentially expressed genes, miRNAs, and siRNAs were detected among and within quinoa varieties during a time-course virus infection study designed to elucidate general response of quinoa and variety-specific interactions to CMV. In addition, sRNASeq analysis was incorporated to complement the transcriptome analysis by characterization of components involved in viral gene silencing. Furthermore, sRNA analysis of quinoa varieties provided knowledge about endogenous and virus-derived siRNAs (vsiRNAs) as well as validated known and novel miRNAs. These findings are of particular interest for molecular breeding of quinoa varieties to enhance resistance against virus.

## 2. Materials and Methods

### 2.1. Plant Materials and Virus Inoculation

Seed from three *Chenopodium quinoa* varieties (Table 1), ‘Jessie’ (sweet), ‘QQ74’ (semi-sweet), and ‘Red Head’ (bitter)] [40, 41] were surface-sterilized in 1% hypochlorite solution (v/v) for 3 minutes followed by three rinses with sterile deionized water. Seed were sown in greenhouse growing media (Promix) and grown in growth chambers under controlled conditions [60% humidity; 16/8 hours (light/dark); 23°C]. Freeze-dried CMV infected tobacco tissues were obtained from ATCC (PV-242). *Nicotiana tabacum* cv T.R.Madole was used as the propagation host. Leaves of seven-week-old quinoa plants were mechanically inoculated with CMV-infected tobacco leaves or mock-inoculated with phosphate buffer (pH:7.0). Inoculated leaves were harvested at 1 or 4 dpi. The experiment was designed as a 3×2×2×3 factorial [three varieties; two treatments (virus or mock); two sample times (1 or 4 dpi); three biological replicates]. Excised leaf samples were immediately frozen in liquid nitrogen and kept at -80°C until RNA extraction. To verify successful inoculation, virus- and mock-inoculated tissues were tested by CMV AgriStrips (Bioreba, Reinach, Switzerland) according to manufacturer’s instruction.

**Table 1.**
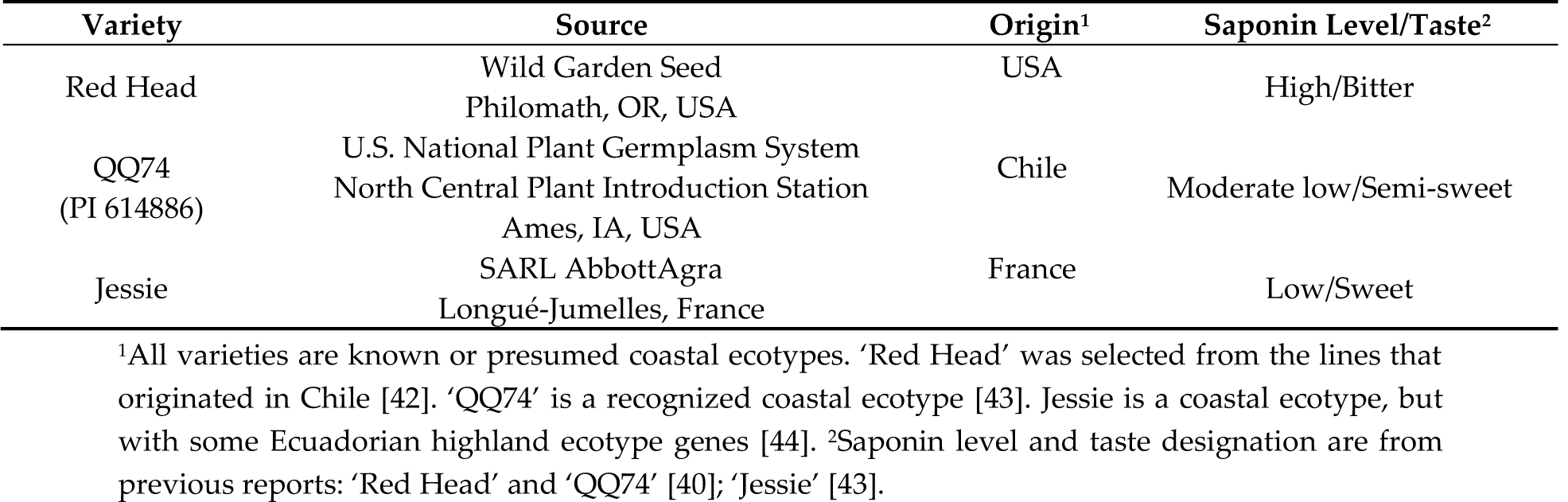
Quinoa (*Chenopodium quinoa*) used in the study.

### 2.2. Total RNA Extraction, Library Preparation, RNA and Small RNA Sequencing

Total RNA was purified from individual samples by Direct-zol RNA MiniPrep Plus kit (Zymoresearch, Irvine, CA, USA) according to manufacturer’s protocol. Concentration of DNA-digested RNAs was measured by NanoDrop™ One Microvolume UV-Vis Spectrophotometer (Thermo Scientific, Waltham, MA, USA). The cDNA library preparation for RNASeq and small RNASeq, and sequencing on illumina HiSeq 4000 was performed by GENEWIZ (South Plainfield, NJ, USA). The same samples were used for RNA and small RNA sequencing.

### 2.3. Genetic Variation Among Quinoa Varieties

The clean reads from transcriptome analysis (see bioinformatics analysis below) were used as input in STAR [45] to map against *C. quinoa* reference genome. Genome Analysis Tool Kit (GATK) [46] was further used to detect SNP and Indels. SnpEff [47] was utilized for functional annotation of variants among quinoa varieties. PHYLIP [48] was utilized to draw phylogenetic trees based on the distance matrix of similarity between variants of the samples.

### 2.4. Bioinformatic Pipelines

The pipeline used to analyze RNASeq data was followed as described previously[49]. In summary, the quality of obtained raw data was checked by FastQC v0.11.5 [50]. Adapters, low quality reads, and sequences shorter than 30 nucleotides were trimmed by Trimmomatic v0.38 [51]. High quality clean reads were mapped to C. quinoa reference transcripts by Salmon v0.12.0 [52] to obtain normalized TPM values. Normalized transcript-level read counts were transformed to gene-level abundance in order to use in differential gene expression R package, DESeq2 [53]. The DEGs were determined in two modes: 1) with a full design model (time×variety×treatment) where DEGs were determined across quinoa varieties (Jessie as reference level) in time-series inoculation (1dpi as reference level) between CMV-inoculated and mock-inoculated samples, and 2) individual variety with a time-specific treatment design (time+treatment+time:treatment), where DEGs were identified in each variety in time-series inoculation (4 dpi vs 1 dpi) between CMV-inoculated and mock- inoculated samples. The DEGs with adjusted *p*-value ≤ 0.05 were considered significant. To identify GO annotation and KEGG pathways of DEGs, KOBAS v3.0 [54] was used. Significant enriched GO terms and KEGG pathways were chosen based on the corrected *p*-value ≤ 0.05 obtained through Benjamini and Hochberg’s method for controlling false discovery rate (FDR).

The pipeline used to analyze sRNASeq data was followed as described previously [55]. Briefly, after inspecting raw reads quality by FastQC, adapters and low-quality bases were trimmed by Trimmomatic. Afterwards, ncRNAs (tRNA, rRNA, snRNA, snoRNA) from NCBI, Rfam, and RepBase databases were removed to obtain clean reads that were employed for downstream analyses. In miRNA analysis, clean reads were mapped to *C. quinoa* reference genome by Bowtie2 [56] to retrieve quinoa sequences. Then, miRDeep2 [57] was employed to detect known miRNAs based on miRBase v22 database, and predict novel miRNAs based on the randfold with a *p*-value < 0.05. Read counts of known and novel miRNAs obtained by “quantifier” module of miRDeep2 were used in DESeq2 for detection of differential expressed miRNAs (DEmiRNA) based on the same design formula as DEG analysis. The DEmiRNA with adjusted *p*-value ≤ 0.05 were considered significant and further analyzed by TargetFinder [58] to predict their target genes. Significant GO terms and KEGG pathways of target genes were obtained by KOBAS as described above. Characterization of endogenous siRNAs and vsiRNAs was carried out by alignment of miRNA-free clean reads to *C. quinoa* reference genome and three RNAs of CMV, respectively, using Bowtie2 with alignment options of very sensitive, zero mismatch, and 18 nucleotides as the minimum sequence size. The 5’ base enrichment and nucleotide size distribution analyses were carried out by “reformat” module of BBMap [59] software. To prepare the read counts for differential expression of siRNA, miRNA-free clean reads were used as input in ShortStack 3.7 [60]; this was used to determine siRNA clusters accumulating in genomic loci. The parameters used in ShortStack were nohp mode, mismatch 0, dicermin 18, dicermax 30, DicerCall cut-off of 80%, and mincov 0.5. Differential expression of siRNAs was determined in two modes as the same as DEG analysis of RNASeq. DE-siRNA clusters from full design and variety-specific models were considered significant based on Benjamini Hochberg adjusted *p*-value ≤ 0.05. To predict putative genes, TE, and TFBS as potential targets of DE-siRNA clusters, a window frame of 2kb upstream of the gene was inspected. Targeted sequences were retrieved and searched using CENSOR [61] algorithm to find candidate TE from RepBase database. Also, the retrieved sequences were subjected to PlantRegMap [62] database against *Arabidopsis thaliana, Beta vulgaris* and *Spinacia oleracea* to perform an exhaustive search on transcription factor binding sites with a *p-value* cut-off of 1e-7. Finally, to annotate the biological function of the targeted gene, KOBAS was used to perform GO and KEGG pathway enrichment analyses.

### 2.5. Validation of DEGs by RT-qPCR

To validate RNASeq results, nine significant DEGs were randomly selected to conduct quantitative RT-PCR on total RNA samples used for RNASeq experiment. To target each gene, two random samples were selected as template: one from mock-inoculated and one from virus-inoculated sample. For each sample, two technical replicates were used. Primers were designed by Primer3 software (Supplementary Table S1). Total RNA was primed by random hexamer and reverse transcribed using SuperScript™ III Reverse Transcriptase (Invitrogen, Carlsbad, CA, USA) according to manufacturer’s instruction. The cDNA synthesis was carried out on Eppendorf Mastercycler Nexus Thermal Cycler (Hamburg, Germany). The PCR reaction was conducted in a final volume of 10µl [1 µl cDNA, 0.4 µl of each primer (10 µM), 3.2 µl RNase/DNase-free water, and 5 µl PowerUp™ SYBR™ Green Master Mix (Applied Biosystems, USA)]. The amplification conditions were 2min at 50°C for UDG activation, initial denaturation of 2min at 95°C followed by 40 cycles of 95°C for 15 s and 60°C for 1min. The qPCR was performed by QuantStudio™ 6 Flex Real-Time PCR machine (Applied Biosystems, Foster City, CA, USA). Elongation Factor 1α (CqEF1α)(XM_021888526.1) was considered the reference gene for normalization of gene expression. Relative gene expression was calculated using the 2^−ΔΔCT^ method [63].

### 2.6. Relative Expression of TSARL1 Gene during Virus Infection

The expression of TSARL1 was relatively quantified upon time-course CMV infection among quinoa varieties. Three biological and two technical replicates was used in RT-qPCR. The primer design (Supplementary Table S1), cDNA synthesis and qPCR methods were performed as described above. The relative expression pattern of the gene was detected based on the comparison of mock- and CMV-inoculated samples using the 2^−ΔΔCT^ method.

### 2.7. Validation of Known and Novel miRNA by Stem-loop RT-PCR

To validate results of detected miRNAs, six known and nine novel miRNAs were randomly selected. Stem-loop primers for cDNA synthesis were designed by miRNA Primer Design Tool [64]. Stem-loop pulsed reverse transcription of each miRNA was according to Varkonyi-Gasic et al. [65] with few modifications. Briefly, 1-2 µg total RNA was added to 0.5 µl dNTP (10mM), 1 µl stem-loop primer (10µM) (Supplementary Table S1) and desired volume of RNase/DNase-free water to a total of 13.8 µl. The mixture was incubated for 5min at 65°C followed by 3 min ice incubation. Subsequently, 4 µl of 5X first-strand reverse transcriptase (RT) buffer mixed with 2 µl DTT and 0.2 µl SuperScript™ III Reverse Transcriptase kit (Invitrogen, USA) (200U/µL) were gently added to yield the final volume of 20 µl. Tubes were loaded to thermal cycler for pulsed reverse transcription with initial incubation at 16°C for 30 min, followed by 60 cycles of 30 °C for 30 s, 42 °C for 30 s and 50 °C for 1 s. Then, the samples were incubated at 85 °C for 5 min to inactivate RT enzyme. To prepare PCR reactions, 1 µl of cDNA was added to 0.4 µl of each primer (10 µM), 8.2 µl water, and 10 µl of Platinum™ II Hot-Start Green PCR Master Mix (2X). Loaded samples in thermocycler were initially denatured at 94°C followed by 35-40 cycles of 94°C for 15s, and 60°C for 30sec. PCR products were resolved electrophoretically on ethidium bromide stained-2% agarose gel immersed in 1× TAE and visualized with Biorad Gel Doc XR instrument (Biorad, USA).

## 3. Results

### 3.1. Symptomatology and Verification of CMV Infection on Quinoa Samples

A total of 36 quinoa samples comprised of three varieties (Table 1), two harvest time-points, and three biological replicates were either inoculated with CMV or inoculation buffer. By 4 days after inoculation with CMV, chlorotic local lesions developed across all replicates of ‘QQ74’ and ‘Jessie’, whereas all replicates of the bitter variety ‘Red Head’ had systemic chlorosis (Figure 1A). Infection was confirmed *in vitro* by immunoassay and *in silico*, with all CMV genomic RNAs detected in RNASeq and small RNASeq datasets (Figure 1B). Control samples that underwent mock-inoculation did not have symptoms or signs of infection.

**Figure 1.**
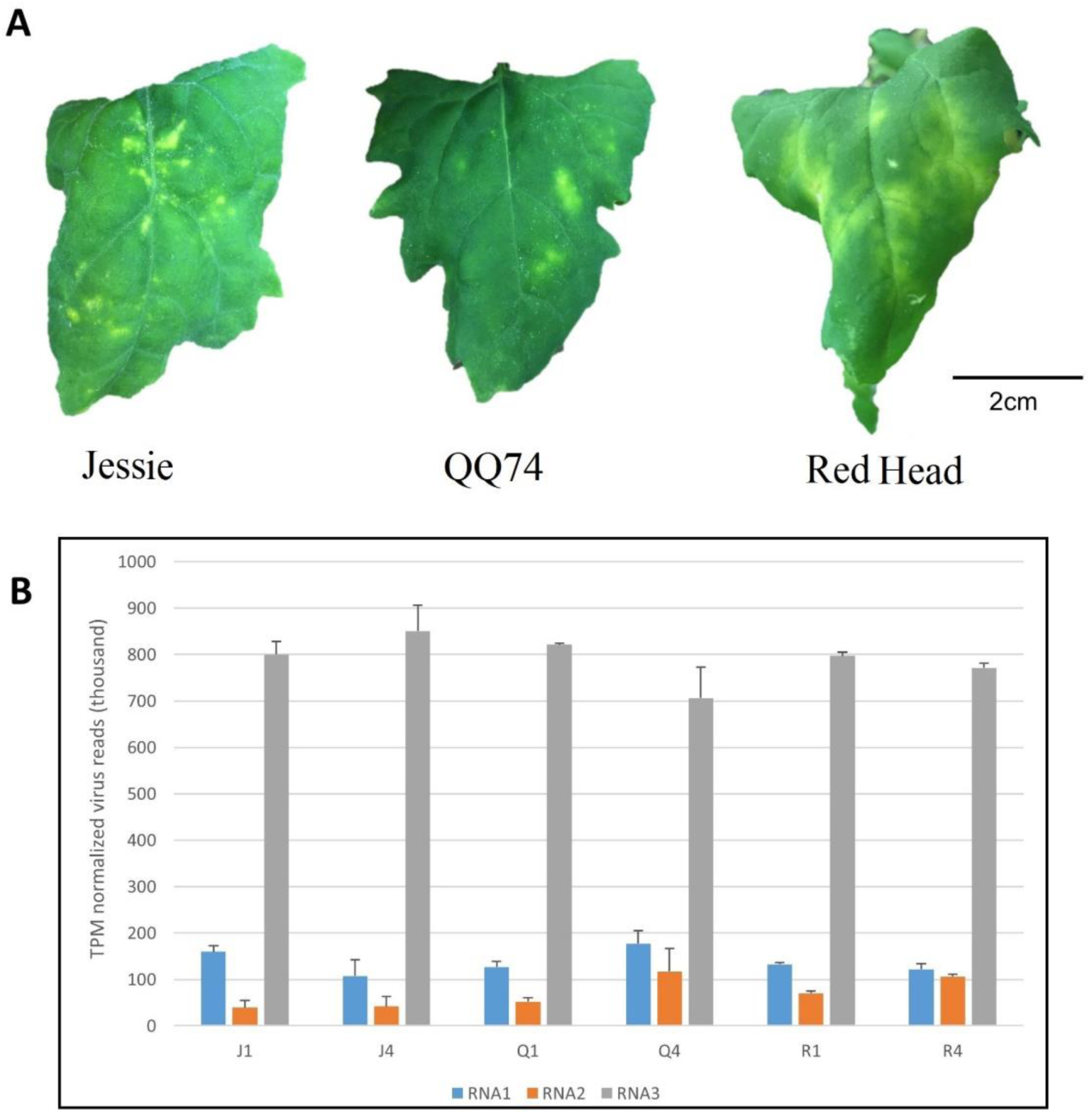
Virus-induced symptoms of *Chenopodium quinoa* varieties and accumulation of viral RNA sequences from deep sequencing data. **(A)** Cucumber mosaic virus (CMV)-induced symptoms on *C. quinoa* varieties at 11 days post inoculation (dpi): Jessie (low saponin level = sweet variety); QQ74 (moderate saponin level = sweet variety); Red Head (high saponin level = bitter variety). **(B)** Normalized (TPM) number of viral reads across CMV RNAs. J, Q, and R are initials of the varieties, and the 1,4 are dpi. The bars indicate standard errors.

### 3.2. Genetic Variation Among Quinoa Varieties

The total of 36 adapter-free, high quality transcriptome data were used in Genome Analysis Tool Kit (GATK)[46] to determine single nucleotide polymorphism (SNP), insertion and deletion (Indel) variation within and between varieties. The sweet variety ‘Jessie’ had higher number of variants with an average of 191,087 in compared to ‘QQ74’ and ‘Red Head’ that averaged 137,252 and 142,407 variants, respectively. There was close relationship among the individual samples of each variety, however they were not identical (Figure 2A). Higher similarity of variants between ‘Red Head’ and ‘QQ74’ than ‘Jessie’ which is genetically distant are highlighted in the phylogram (Figure 2B).

**Figure 2.**
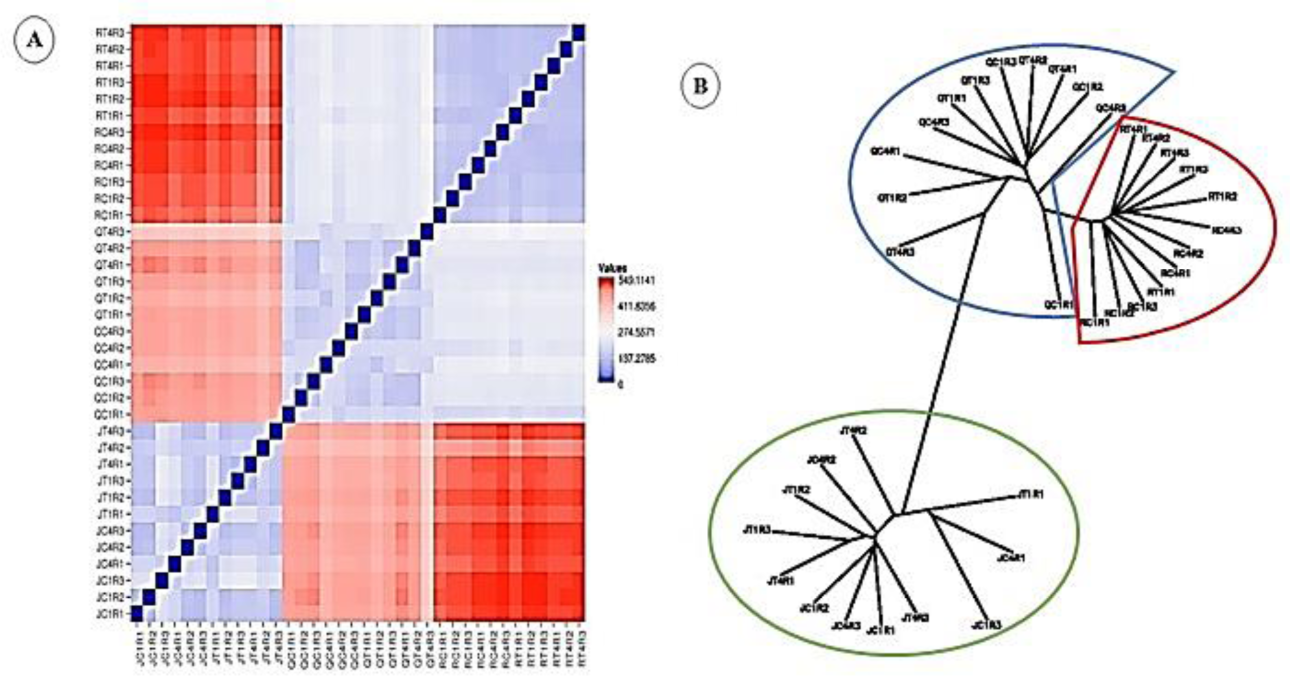
Genetic variation analysis of quinoa varieties. **(A)** Distance matrix within and among samples. In color legend, the blue representing the closest distance between samples and the red represents farther distance between samples. **(B)** Unrooted phylogenetic tree of quinoa varieties with their biological replicates. J, Q, and R are initials of the varieties, the 1,4 are dpi, and respective biological replicates represented by either R1, R2, or R3.

### 3.3. Transcriptome and Small RNASeq Profile of Quinoa Samples

To determine molecular interactions and stress responsive mechanisms in quinoa during CMV infection, transcriptome and small RNA sequencing were performed. In transcriptome sequencing, a total of 1.182 billion raw reads was generated. After removing low quality reads and adapters, trimmed sequences (1.131 B) were mapped to *C. quinoa* reference transcripts [4] with an average successful alignment percentage rate of 90.17 (Supplementary Table S2). To eliminate the effect of different sequence length and depth of read coverage among sample libraries, the mapped reads were normalized to Transcript Per Kilobase Million (TPM) values for use in downstream analyses.

The small RNA sequencing of 36 samples resulted in 364.31 million raw reads. After the trimming step, the surviving reads (327.84 M) were mapped to *C. quinoa* reference genome with an average successful alignment percentage rate of 97.93 (Supplementary Table S2). Non-coding (nc)RNAs including tRNA, rRNA, snRNA, snoRNA were subsequently removed from mapped reads, and the remaining reads (198.11 M) were further used for miRNA or siRNA analyses.

### 3.4. Differential Expressed Genes (DEG) Identification and GO and KEGG Pathway Enrichment Analyses

TPM normalized gene counts were imported in DESeq2 package, and DEGs were determined between CMV- and mock-inoculated samples with an adjusted *p*-value cut-off ≤ 0.05. In the first mode of analysis (full design) that encompassed the effect of variety, time and treatment and their interactions, a total of 250 genes were differentially expressed (Table 2, Supplementary Table S3). Principal component analysis (PCA) was used to compare gene expression profiles of the samples; samples with similar pattern were plotted close to each other while samples with different profiles scattered far from each other in the scores plot of the PCA (Supplementary Figure S1). PC1 explained 53% of variation and PC2 explained 12%. In the variety-specific analysis (individual variety design), DEGs were determined between CMV-inoculated and mock-inoculated samples by including time, treatment and time × treatment. In ‘Jessie’ a total of 332 genes, in ‘QQ74’ 85 genes, and in ‘Red Head’ 140 genes were differentially expressed (Table 2).

**Table 2.** Total numbers of DEGs, enriched GO terms and KEGG pathways (*p* ≤ 0.05) assigned to DEGs of *Chenopodium quinoa* varieties upon cucumber mosaic virus infection.

| Design | Full model |  | Variety-specific model |  |  |  |  |  |
| --- | --- | --- | --- | --- | --- | --- | --- | --- |
|  | All varieties |  | Jessie |  | QQ74 |  | Red Head |  |
| Expression pattern | UR | DR | UR | DR | UR | DR | UR | DR |
| DEGs | 142 | 108 | 150 | 182 | 31 | 54 | 73 | 67 |
| GO terms | 134 | 223 | 217 | 325 | 119 | 248 | 234 | 451 |
| KEGG pathways | 9 | 1 | 4 | 7 | 8 | 15 | 4 | 7 |

To identify biological function of DEGs, Gene Ontology (GO) term analysis was performed on DEGs using KOBAS with corrected *p*-value cut-off ≤ 0.05 (Table 2). In the full design, top significant up-regulated (UR) GO terms were endopeptidase regulator and oxidoreductase activities, and the top terms for down-regulated (DR) GO terms were cell aging and hydrolase activity. In individual variety design, for ‘Jessie’ top significant UR GO terms were response to oxygen-containing compound and flavonoid biosynthetic process, whereas photoperiodism and response to stimulus were among top DR GO terms. In ‘QQ74’, cellulose catabolic process and telomere maintenance were top two significant UR GO terms, and telomere organization and beta-glucan catabolic process were top DR GO terms. In ‘Red Head’, top UR GO terms were chloroplast accumulation movement and response to stress, and top DR GO terms were response to chemical and chlorophyll metabolic process (Supplementary Table S4). To identify the role of DEGs in primary and secondary biological pathways, KEGG enrichment analysis was performed with KOBAS (Table 2). With all the experimental factors in the statistical design, glycerophospholipid metabolism was the top significant UR KEGG pathways; and pyrimidine metabolism was the only enriched DR KEGG pathway. Within each variety design, in ‘Jessie’, flavonoid biosynthesis was the most enriched UR KEGG pathways, and circadian rhythm was top significant DR KEGG pathways. In ‘QQ74’, top enriched UR KEGG pathway was sulfur metabolism, whereas the most significant DR pathway was biosynthesis of secondary metabolites. In ‘Red Head’, top enriched UR KEGG pathway was circadian rhythm, and the most significant DR pathway was glycerolipid metabolism (Supplementary Table S4). Among the enriched GO terms and KEGG pathways, modulation of hormonal signaling, plant-pathogen interaction (PPI), photosynthesis, carbon metabolism, and amino acid biosynthesis were shared at least once between varieties.

### 3.5. Differential Expression of TSARL1 gene in Quinoa upon CMV Infection

To inspect the effect of local and systemic infection of CMV on expression of TSARL1 gene in sweet/semi-sweet and bitter quinoa varieties, respectively, mock- and virus-inoculated samples were compared by RT-qPCR. In sweet varieties, TSARL1 expression was consistently upregulated at 1 and 4 days post inoculation (dpi). However, in the bitter variety gene expression was upregulated at 1dpi but suppressed drastically at 4dpi (Supplementary Figure S2). Therefore, there was a concomitant decrease in expression of TSARL1 and systemic infection of CMV in the bitter variety. However, in sweet varieties, elevation of TSARL1 expression and local lesions were concurrent.

### 3.6. Detection, Differential Expression, and Target Gene Analysis of Known and Novel miRNAs

To further characterize miRNAs from trimmed small RNASeq reads, sequences mapped to *C. quinoa* were separated from ncRNA (tRNA, rRNA, snRNA, snoRNA), and the resulting clean reads were used as input for miRDeep2 software. Known miRNAs were detected based on the mature and precursor sequences of miRBase v22. The total of 81 known miRNAs belonging to 11 miR families (miR 156, 160, 162, 166, 171, 172, 319, 393, 395, 398, and 399) were detected, with miR166 as the most abundant detected known mature miRNA among the quinoa samples. In addition, novel miRNAs were identified based on the randfold (*p* < 0.05) and miRDeep2 score ≥ 0. Prediction of novel miRNAs resulted in 876 significant novel mature miRNAs, among which novel miRNA “NW_018742204.1_181” was the most prevalent novel miRNA among quinoa samples (Supplementary Table S5).

Determination of RISC factors involved in miRNA biogenesis is possible through nucleotide size distribution and 5’ base enrichment analyses of miRNAs. The dominant size and the enriched 5’ terminal base of known miRNAs were cleaved sequences with the size of 21 nucleotides and 5’ Uracil (U) (Supplementary Table S5). To characterize differential expressed miRNA (DEmiRNA), read counts of known and novel miRNAs were used in DESeq2, and DEmiRNAs were determined between CMV-inoculated and mock-inoculated samples based on the threshold *p* ≤ 0.05. In the full model that included the variety interaction as well, one DEmiRNA was UR and one was DR. Within the varieties, in ‘Jessie’ two UR- and four DR-miRNAs (including miR166, 399) were detected, in ‘QQ74’ five DEmiRNAs (5 DR) were identified, and in ‘Red Head’ no differentially expressed miRNA was detected. TargetFinder analysis of miRNAs detected a total of 126 candidate target genes that four (3 UR, 1 DR) belonged to the full-model design and 122 (19 UR, 103 DR) belonged to individual- variety design (Table 3, Supplementary Table S6). To provide comprehensive function of DEmiRNAs, enrichment analysis of targeted genes was performed by KOBAS (adjusted *p* ≤ 0.05). Overall enriched biological pathways were involved in cell recognition and water homeostasis. In ‘Jessie’, targeted genes were enriched in ether lipid/glycerophospholipid metabolism, ubiquitin-mediated proteolysis and endocytosis functions. In ‘QQ74’, no biological pathway was enriched (Supplementary Table S6).

**Table 3.**
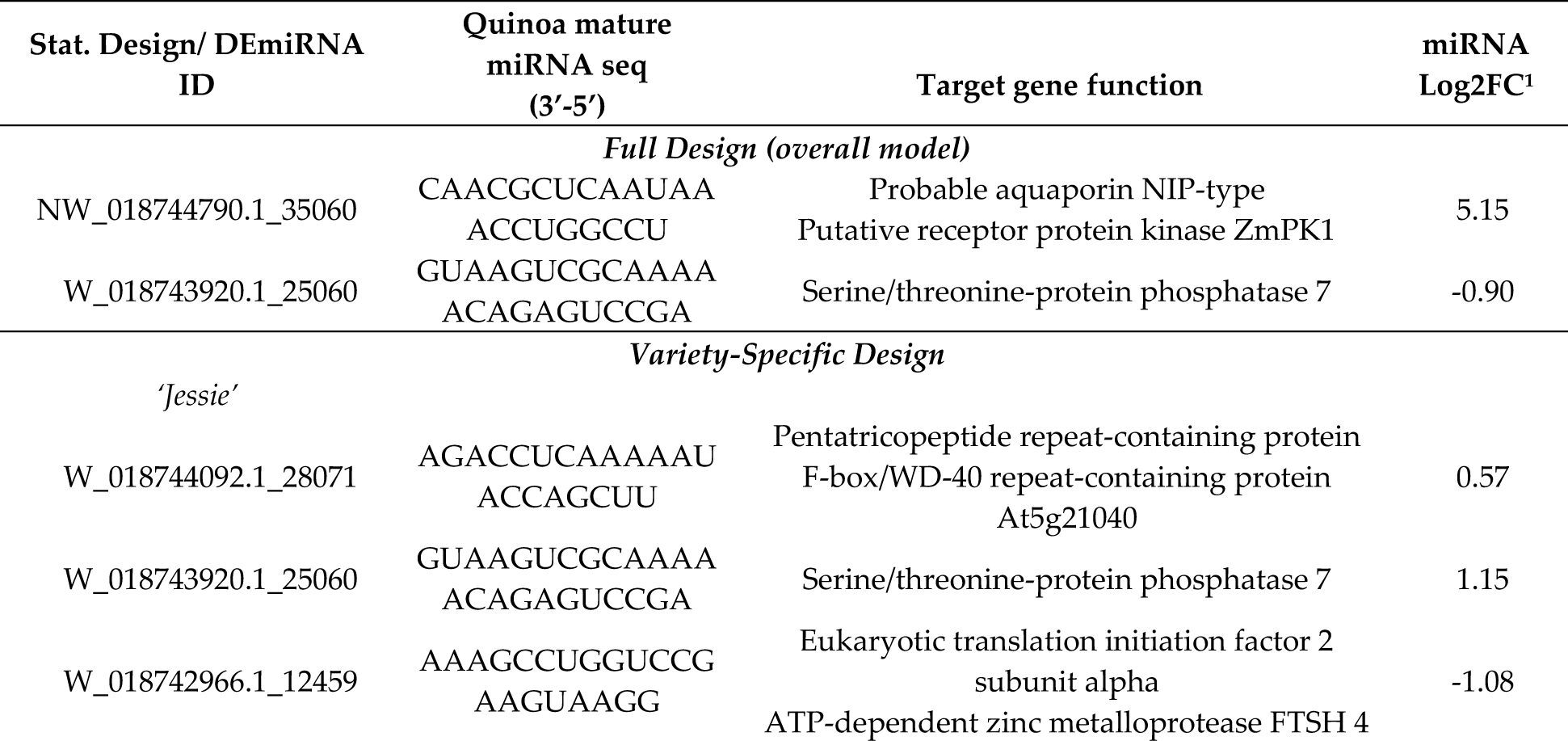
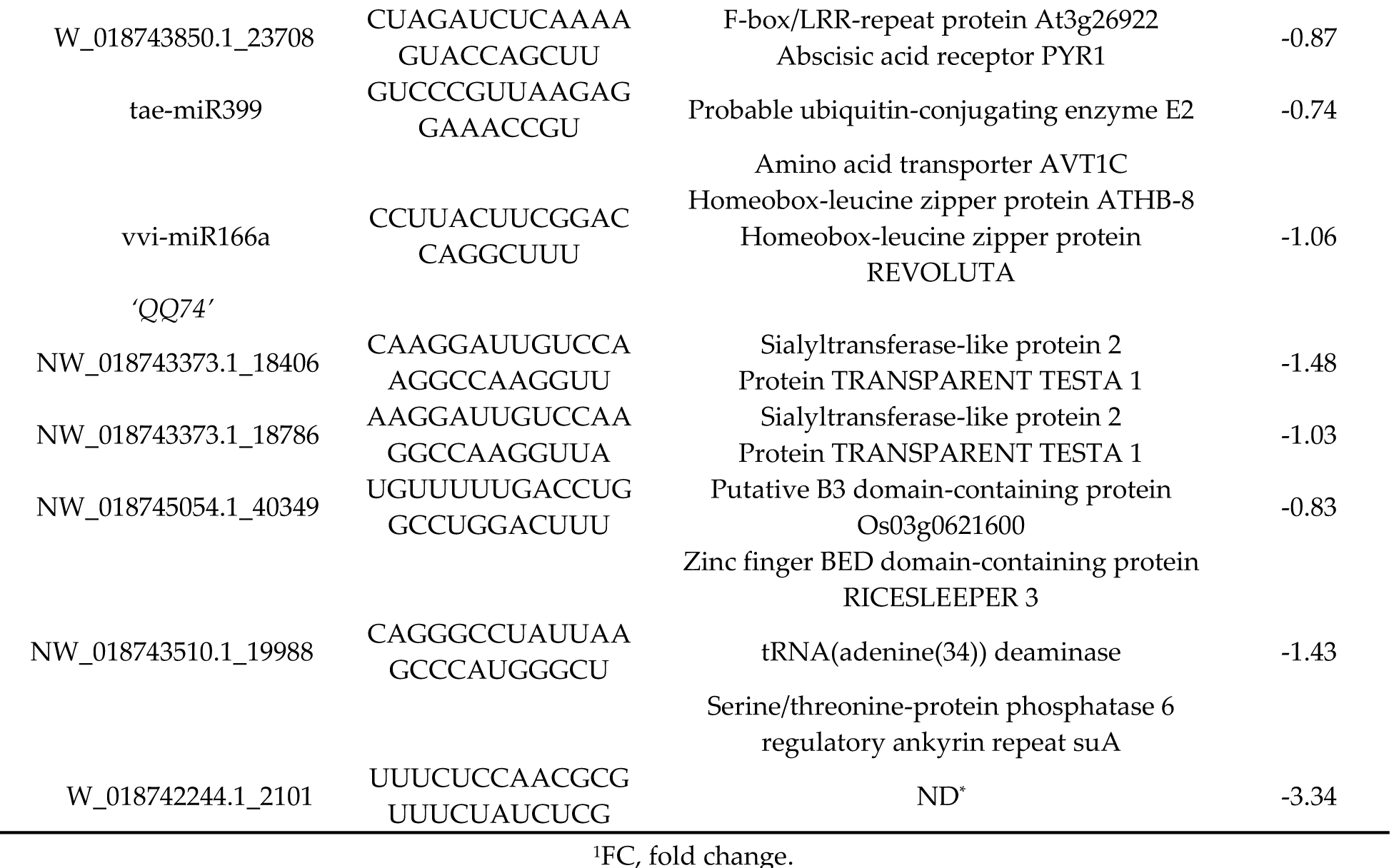
Differential expressed known and novel microRNAs (DEmiRNA) between cucumber mosaic virus-inoculated and mock-inoculated samples among and within quinoa varieties with their top two candidate target genes.

To identify the complexes involved in formation of endogenous siRNA, nucleotide size distribution and 5’ end base abundance of detected siRNAs are needed. Since siRNAs completely match their target sequences, mapping clean small RNA reads to *C. quinoa* reference genome allowing zero mismatch resulted in siRNA targets. These target sequences were used to identify dominant cleaved nucleotide size and 5’ terminal base of endogenous siRNAs. The average alignment rate was 99.75%. The sequences with 24 nt were more abundant than other nucleotide sizes (Figure 3). Also, Adenine (A) was the predominant base at 5’ end (Supplementary Figure S3). To enable differential expression of siRNAs, the ShortStack package was used to determine the counts of siRNA clusters varying in size (18-30 nt) over quinoa genome. A total of 160,533 siRNA clusters was retrieved per individual sample (Supplementary Table S7). The abundance of overall siRNA sizes varied among the varieties, harvest time and treatments; however, the abundance of 24nt clusters were significantly (*p* = 0.01) higher in ‘Red Head’, bitter variety, than two sweet/semi-sweet varieties, ‘Jessie’ and ‘QQ74’ (Figure 3). Using DESeq2 on cluster counts of the samples by comparing CMV-inoculated and mock-inoculated tissues under two models, overall analysis and variety-specific analysis, revealed that the size of 24 nt was the most abundant differentially expressed Dicer cleaved siRNA. Among the total 377 differentially expressed siRNA clusters, 235 belonged to overall analysis, and 142 belonged to variety analysis (‘Jessie’: 100; ‘QQ74’: 39; ‘Red Head’: 3) (Supplementary Table S8). These 24nt hc-siRNA clusters were used for further analyses. Since hc-siRNAs target genes and intergenic regions such as transposable elements, gene promoter and transcription factor binding sites, multiple methods were exploited to predict candidate targets spanning 2kb of the targeted genes. Based on the overall or variety-specific analyses, targeted repetitive regions and biological functions of the targeted genes were annotated separately (Supplementary Table S9).

**Figure 3.**
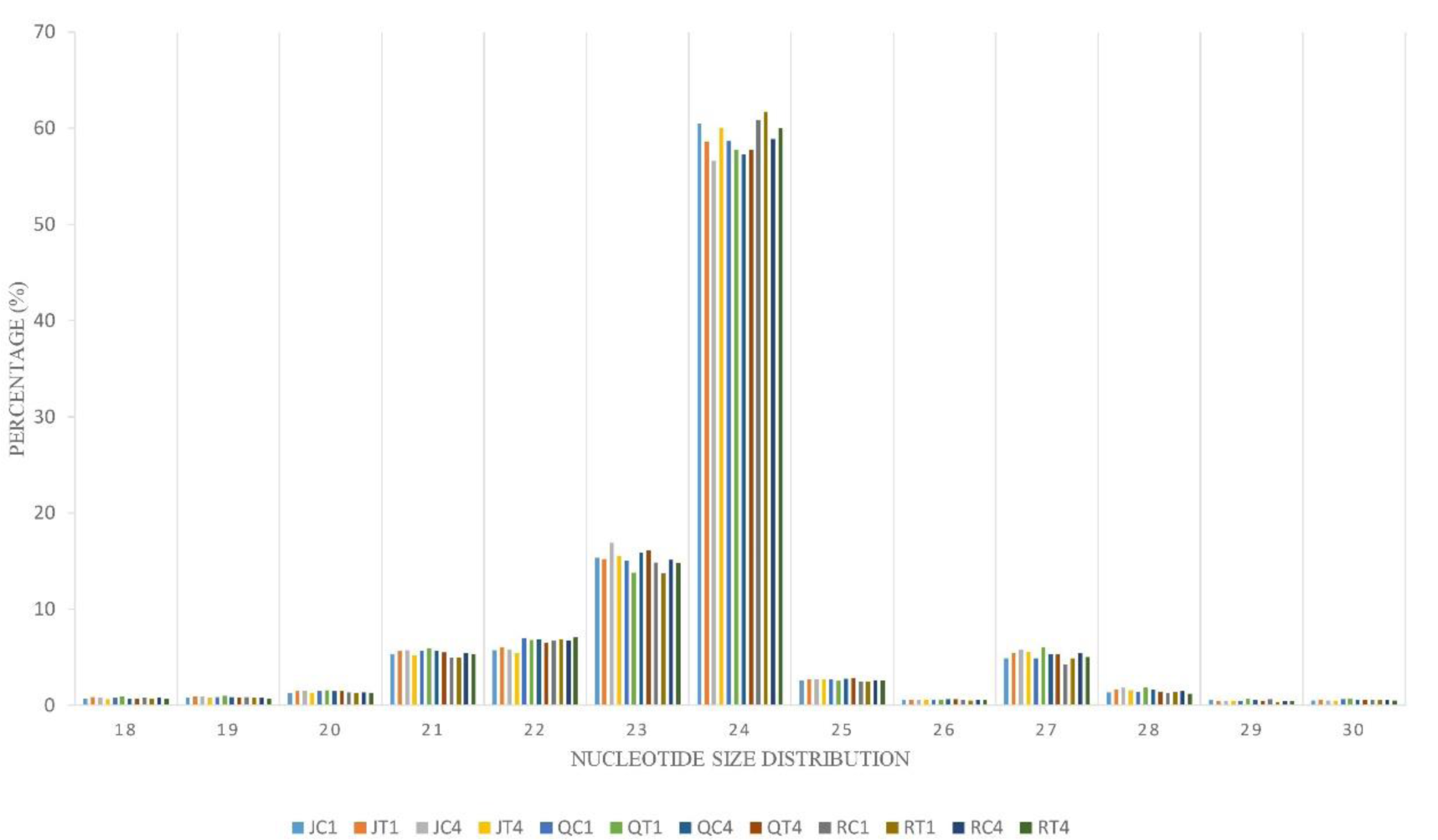
Nucleotide size distribution of endogenous siRNA. This graph depicts data from quinoa varieties (J, Q, R) with two treatments (C,T) at two time periods (days after inoculation [1,4]). J, Q, and R are initials of the varieties, and C and T are mock- and virus-inoculated, respectively.

### 3.8. Characterization of Virus-Derived siRNAs

Complexes involved in RNAi leading to antiviral defense could be identified through mapping clean reads to CMV genome with the perfect match option. Therefore, the same alignment option as described above was followed to retrieve vsiRNA sequences. These sequences provide information about nucleotide size distribution and 5’ end base enrichment of vsiRNA that have been cleaved by plant RNA silencing factors. Forty five percent of vsiRNA had a length of 21 nt, and 30% were 22 nt (Figure 4A). There was no consistent 5’ terminal base among varieties and time points (Figure 4B, Supplementary Figure S4). Obtained vsiRNA sequences also provided information about three RNA segments of CMV from which they were derived. Sequences derived from viral RNA3 were more prevalent than signatures of RNA1 and RNA2 (Figure 5).

**Figure 4.**
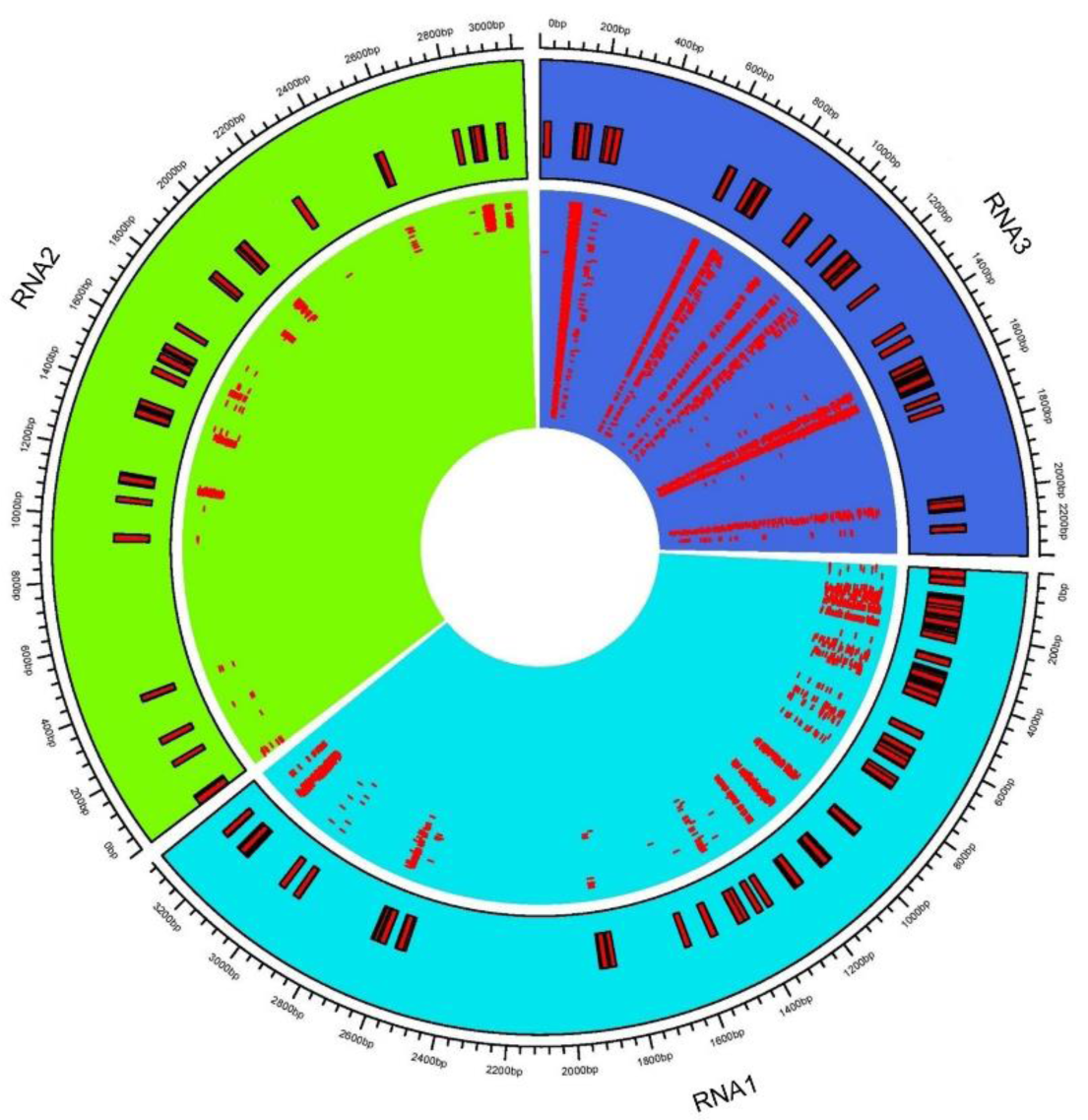
Cucumber mosaic virus (CMV)-derived small interfering RNA (vsiRNA) information. (A) Nucleotide size distribution of vsiRNA. (B) Enrichment of 5’ terminal base across the varieties and days post inoculation. The graph depicts data from quinoa varieties (J,Q,R), and time (days after inoculation).

**Figure 5.**
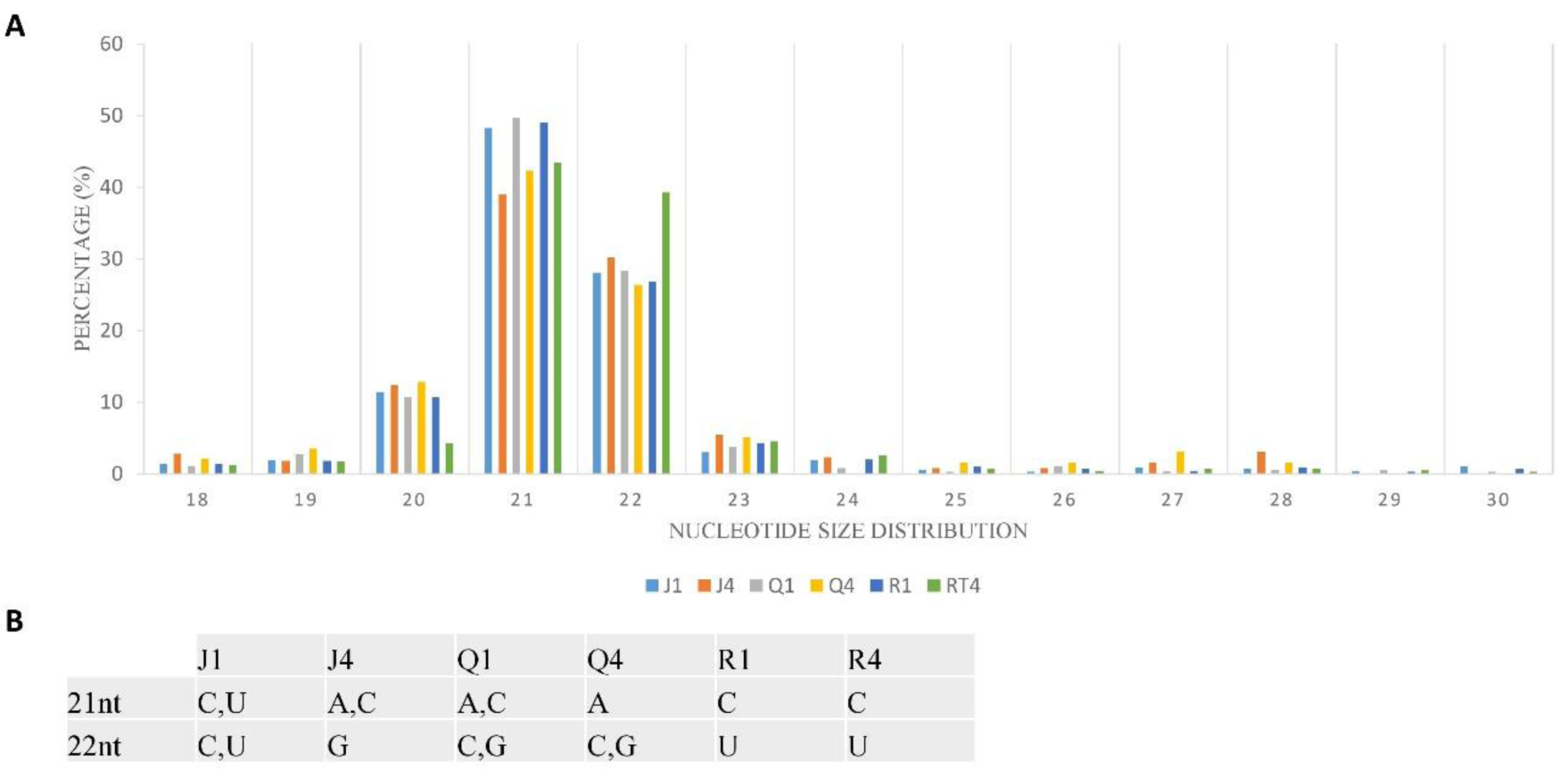
Distribution of virus-derived siRNA (vsiRNA) across cucumber mosaic virus (CMV) RNAs. The consensus vsiRNA distribution across three CMV RNA1, RNA2, and RNA3. The inner rectangles show reads mapped to respective viral RNA. Outer rectangles relate genome coordinates of consensus reads mapped to CMV RNAs.

### 3.9. RT-qPCR validation of DEGs

RT-qPCR with specific primers was used to validate DEG patterns detected *in silico* from RNASeq for nine genes involved in RNA transport, membrane structure, lipid or protein biosynthesis. The resultant fold changes between CMV- and mock-inoculated samples were compared to those of detected DEGs from DESeq2 output. All tested genes had the same expression patterns (Figure 6).

**Figure 6.**
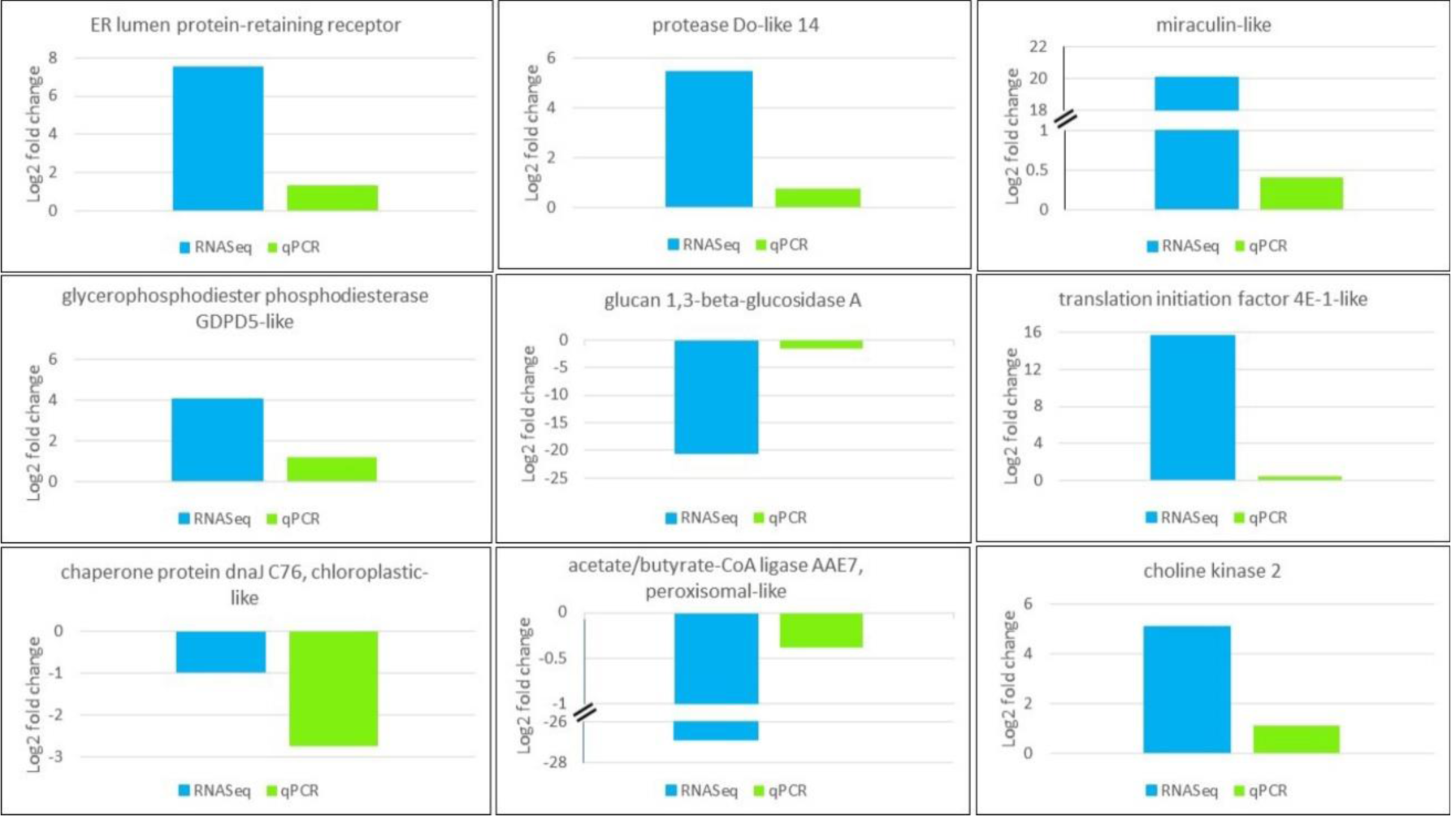
Validation of expression pattern of genes selected from RNASeq (blue) using RT-qPCR (green). CqEF1α was used for gene normalization purpose. Gene function is shown on top of the plot.

### 3.10. Validation of Known and Novel miRNA by Stem-loop RT-PCR

To confirm identified known and novel miRNA *in vitro*, 6 randomly selected known miRNAs including miRs 156, 166b, 166m, 393, 395, 399, plus the 9 novel miRNAs were reverse transcribed using stem-loop primers, followed by PCR and gel visualization. All the tested known and predicted miRNAs were amplified as expected (Figure 7), indicating reliability of the obtained sRNASeq results.

**Figure 7.**
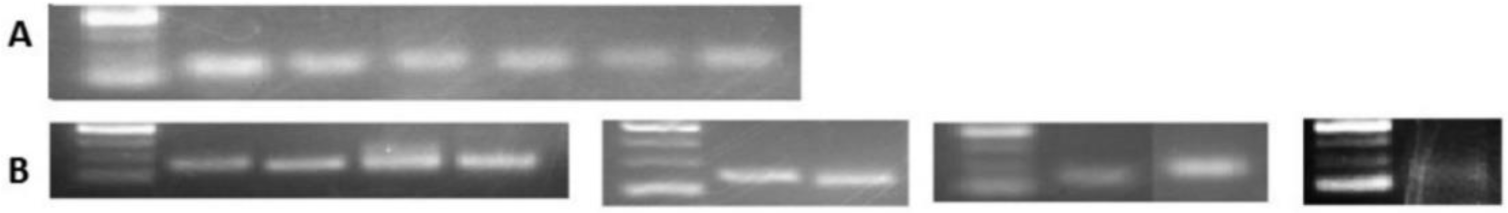
Validation of detected known and predicted novel miRNAs by PCR. From left to right the samples are as follows: **(A)** Known miRNAs 1) ladder, 2) 166b, 3) 166m, 4) 393, 5) 395, 6) 156, 7) 399. **(B)** Novel miRNAs 1) ladder, 2) n2, 3) n5, 4) n6, 5) n9, 6) ladder, 7) n3, 8) n4, 9) ladder, 10) n1, 11) n7, 12) ladder, 13) n8. The ladder is O’range Ruler 50bp ThermoFisher with the range of 50 to 200 with 50bp increments.

## 4. Discussion

Transcriptional responses of different quinoa ecotypes during abiotic stresses [66, 67] and transcriptome changes of herbaceous host, *C. amaranticolor*, in response to different viruses [36-38] have been documented. However, the global gene expression pattern and small RNA profile of quinoa varieties varying in saponin level during virus infection was lacking prior to this study. To inspect the molecular interaction of CMV infection on the host transcriptome over time (1 and 4 dpi), three varieties with low to high seed saponin content were inoculated with either buffer or CMV-infected leaf sap. Based on bioinformatic analysis of transcriptome sequences, DEGs were detected between mock- and virus-inoculated samples. Furthermore, integration of sRNA analyses including miRNA, endogenous siRNA, and vsiRNA provided the knowledge on complexes recruited in host sRNA biogenesis and filled the knowledge gaps on antiviral gene silencing factors in quinoa during CMV infection.

Genetic variations and gene expression clustering among and within quinoa varieties in this study were evaluated through multiple methods to determine genetic relationships of these quinoa varieties that vary in seed saponin content. The analysis among the varieties generated three clusters, showing highest genetic distance and variation between ‘Red Head’ and ‘Jessie’, varieties for which high and low level of seed saponin have been reported, respectively. The smaller genetic distance between ‘Red Head’ and ‘QQ74’ may be the result of their shared Chilean ancestry. ‘Red Head’ was derived from Chilean landraces, and ‘QQ74’ was collected in Chile. Both are coastal ecotypes. Like most European varieties [68], ‘Jessie’ is primarily of Chilean coastal ancestry [68] but also has Ecuadorian highland ecotype in its genetic background [44]. Within the varieties, clustering of individual samples indicated close genetic distance and relationship, and similarity of biological replicates regardless of the treatment or harvesting time, however, they were not identical.

After CMV infection, virus-induced symptoms on ‘Red Head’ spread systemically, whereas in sweet/semi-sweet varieties response was limited to local chlorotic lesion. This negative correlation between virus spread and relative saponin content of quinoa seed is consistent with a previous study [69]. In that study, tomato yellow leaf curl China virus (TYLCCNV)-infected plants had decreased terpenoid expression, and this led to higher fitness of *Bemisia tabaci*, an aphid vector of TYLCCNV. On the other hand, virus-free plants showed higher expression of terpenoids, which resulted in low level of insect infestation. In quinoa, expression of TSARL1 gene is correlated with saponin content [4]. In this study there was a concomitant decrease in expression of TSARL1 and the systemic spread of CMV in the bitter variety. Since CMV can be transmitted by vectors, it is possible that virus infection increases fitness of the vector in quinoa.

Several metabolic pathways including purine and pyrimidine metabolism, nitrogen processes, and terpenoid backbone were downregulated in virus-infected plants. Purine catabolism is a vital housekeeping function of the plants for growth, development, nitrogen remobilization, and induction of defense-related hormone signaling [70, 71]. Allantoin is a nitrogen-rich compound and an essential intermediate in purine catabolism that typically accumulates in stressed plants [71]. Allantoin induces jasmonic acid (JA) [70, 71], the defense signaling phytohormone that results in systemic resistance against viruses [72-75]. In this study, downregulation of allantoin and nitrogen compounds could have resulted from suppression of purine catabolic processes. These successive suppressions may have decreased JA signaling resulting in plant susceptibility to virus infection [72, 75]. Based on retrieved vsiRNAs from sRNA analysis in our study, there were few sequences of 3’ end of RNA2 encoding 2b protein of CMV. Because viral 2b protein inhibits cleavage activity of Argonaute [76], increased levels of this protein allow the virus to counter both plant defense through JA signaling repression and facilitates long distance movement of the virus. Purine and pyrimidine together as part of nucleotide metabolism of host are essential for virus replication. Downregulation of these two compounds could be an antiviral mechanism that deprives virus of nucleotides required for virus replication machinery [77]. Terpenoids are secondary metabolites with the anti-herbivory and defensive roles in plants [13]. Downregulation of DEG involved in terpenoid backbone biosynthesis was related to a chloroplastic terpenoid pathway (MEP) (Table 4); this restricts biosynthesis of downstream terpenoid compounds and impairs the terpenoid-related defensive roles [69, 78].

**Table 4.**
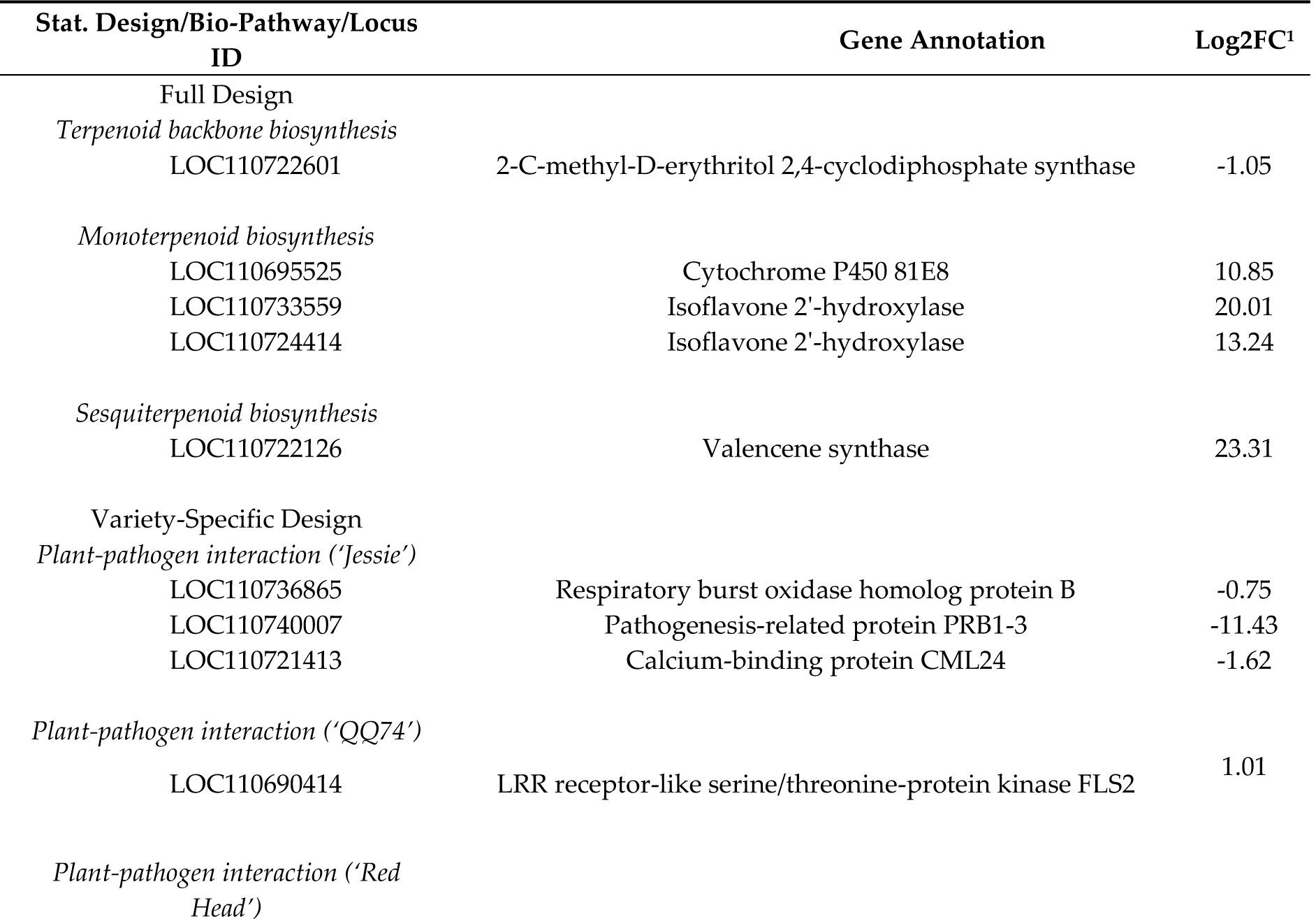
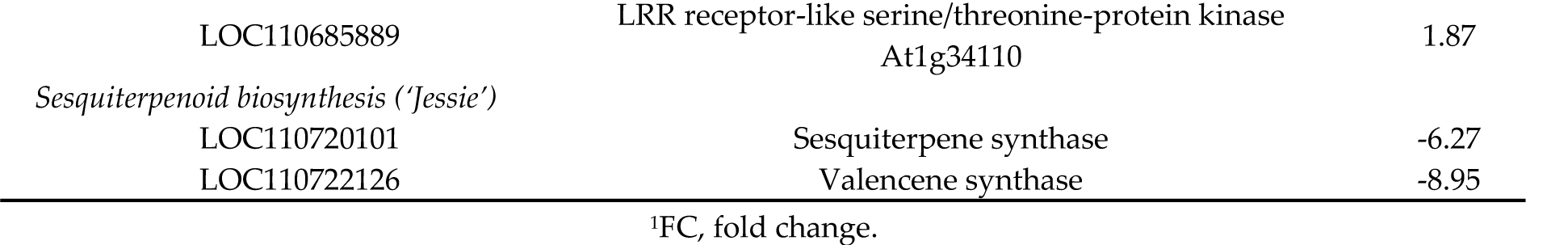
Annotation of detected DEGs between cucumber mosaic virus- and mock-inoculated samples involved in terpenoid biosynthesis and plant-pathogen interaction pathways.

Upregulated DEGs were involved in lipid and nitrogen metabolism, translation, hormone signaling, protein and terpenoid biosynthesis. Nitrogen is one of the primary limiting factors for plant growth, and its assimilation is required for biosynthesis of amino acids glutamate, glutamine, asparagine, and aspartate [79, 80]. CMV infection in tobacco leaves led to overexpression of two enzymes implicated in primary assimilation of nitrogen, glutamate dehydrogenase (GDH) and cytosolic glutamine synthetase (GS1) [81]. In our study, accumulation of nitrogen compounds likely led to induction of amino acid biosynthesis, thus resulting in elevated levels of translation process and protein formation; the finding is also consistent with the study on *Arabidopsis thaliana* that reported impact of elevated level of nitrogen on increasing protein content [79]. Endoplasmic reticulum (ER), the replication site for viruses in the *Bromoviridae* family, is the site for formation of viral factories [82, 83]. In quinoa, following induction of DEGs in nitrogen and protein biosynthesis, amino acid transport to ER was also increased. Because amino acid transport is correlated to abundance of nitrogen in plant cells [80], elevated processes of protein retention in ER and translation might be due to viral demand for the protein biosynthesis required for its replication and assembly. Also, increased level of lipid metabolism might be the result of recruiting host cellular factors to synthesize glycerophospholipids, which is the main component of membraneous viral factories [84], vesicles that are required for virus replication. Phytohormones have several regulatory roles in plant development, growth, host-pathogen interaction, and plant defense mechanism [85]. Salicylic acid (SA) is one of the major defense-related hormones and plays a positive regulatory role against plant viruses. The SA interaction with brassinosteroid (BR) or ethylene (ET), as two other phytohormones, results in either synergistic or antagonistic immune responses. Because BR induces ET biosynthesis [86] and ET precedes the SA signaling pathway [87], the upregulation of both BR and ET DEGs in this study, that was concurrent with overexpression of SA-related DEGs, could have been a synergistic defense response against virus. Enzymatic reactions of synthases, reductases, kinases, etc. in the terpenoid backbone biosynthesis concatenates isoprene units to produce various classes of terpenoids. Monoterpenoids are biosynthesized through MEP pathway, whereas sesquiterpenes are produced in a cytosolic pathway (mevalonate [MVA]) [88]. Overexpression of DEGs pertaining to sesquiterpenes (Table 4) in quinoa might have resulted from increased activity of MVA pathway, which probably compensates the suppressed sources of chloroplastic-derived isoprene units. Although, MEP pathway was downregulated, induction of monoterpenoid DEGs might be from elevated isoprene blocks originated from MVA pathway. The cross-talk of MEP and MVA pathways have been reported in northern red oak as a response to ozone [89].

Analysis on the effect of CMV infection over time on separate quinoa varieties resulted in identification of DEGs that modulated variety-specific biological pathways, such as plant-pathogen interaction (PPI), DNA replication, mismatch and nucleotide excision repair, homologous recombination, and hormone signaling. The DEGs for PPI include defense-related mechanisms that involve reactive oxygen species (ROS) and result in hypersensitive response, cell programmed death, and induction of genes that are related to defense against viruses. All these responses modulate levels of defense hormones such as SA, JA, and ET [90]. In ‘Jessie’, a variety with CMV-induced local lesions, downregulation of PPI was concomitant with suppression of DEGs of ROS and defense hormones (SA, JA, ET), which implies over time (from 1 dpi to 4 dpi), these defense responses were suppressed. Although the fact that a local lesion variety has downregulated defense system seems counter- intuitive, plant reaction to viral inoculation occurs rapidly and processes that require upregulation can be completed within the first hours after inoculation [91]. In ‘Red Head’, the variety with CMV-induced systemic symptoms, both PPI and ROS were upregulated, which implies elevation of these two stress responsive mechanisms over time. Therefore, in contrast to ‘Red Head’ (susceptible host) that had a delayed defense response, ‘Jessie’ (resistant host) expressed defensive pathways rapidly at early stage of infection. Replication and repair processes of host genome is a critical and substantial antiviral mechanism [90]. In tobacco infected with TMV, upregulation of plant DNA homologous recombination resulted in persistent induction of DNA rearrangement and leucine-rich repeat (LRR) gene expression, which led to persistent and broad-spectrum resistance to TMV infection in tobacco [92, 93]. In ‘QQ74’, a variety with few CMV-induced local lesions, PPI over time was overexpressed. This finding could be due to upregulation of DNA replication, mismatch and nucleotide excision repair, and homologous recombination that provided prolonged and persistent defense responses such as PPI even at 4dpi. The DEGs involved in hormone signaling were altered in ‘Jessie’ and ‘Red Head’ but not QQ74. Reprogramming of auxin biosynthesis and/or signaling through altering its function and subcellular localization by several plant viruses including CMV has been shown to be beneficial for virus movement and replication, as well as enhanced infection and disease symptoms [85, 94]. Elevated levels of auxin repressed the hypersensitive response (HR) resulting from SA signaling pathway [85]. In this study, ‘Jessie’ and ‘Red Head’ displayed induction of genes of auxin signaling over time in CMV infection, suggesting suppression of systemic resistance in both varieties via disruption of SA-mediated defense responses. It has been reported that in cucumber infected with CMV, ET is a determinant for viral-induced symptoms [95]. In ‘Red Head’, overexpression of ET-related genes at 4 dpi could be responsible for appearance of CMV-induced systemic symptoms. Abscisic acid (ABA), a phytohormone involved in developmental processes, such as seed germination, fruit ripening, and responses to abiotic stresses [85], is antagonist to SA/JA-dependent defense pathway, ROS production and HR, and therefore represses systemic resistance against viruses [85, 96]. Furthermore, ABA induces callus deposition in plasmodesmata, which limits virus intercellular movement, and also elevates AGO1 activity implicated in gene silencing [85, 96]. In this study, in ‘Red Head’, induction of ABA responding genes at 4 dpi may have played a dual role against CMV infection: 1) utilization of ABA by virus to negate host systemic resistance, and 2) induction of ABA by host to repress virus through limiting viral movement or silencing viral genes via RNAi machinery. In tobacco, cytokinin (CK) accumulation led to enhanced viral resistance through SA-mediated defense responses [97]. In ‘Jessie’ and ‘Red Head’, downregulation of CK-related genes over time was likely related to suppression of systemic resistance through SA signaling pathway. In ‘Jessie’ this might be a host strategy to bring hormonal levels to pre-infection levels, because the virus has already been deactivated in the adjacent cells due to appeared local lesions [98]. Conversely, in ‘Red Head’, suppression of CK-related genes was probably governed by CMV, because of the systemic infection of the virus that led to continuous modulation of defense hormone signaling over time.

The primary small plant-derived RNAs (miRNA and siRNA) are necessary regulatory factors for modification of endogenous or exogenous gene expression through either mRNA cleavage or translation inhibition. RISC factors such as DCL and AGO have crucial roles in miRNA and siRNA biogenesis. Since, in other plants DCL and AGO factors have been identified based on cleaved nucleotide size and 5’ terminal base of sRNAs [24, 27, 28], in this study we conclude that miRNA sizes of 21 nucleotides were cleaved by DCL1, and the single stranded RNAs with 5’U are loaded on AGO1 to guide the RISC to regulate the target genes. In *Arabidopsis*, 24nt siRNAs were cleaved by DCL3 and loaded on AGO4/6/9 to activate genomic and intergenic regions through sequence methylation and histone modification. RDR2 is also recruited in the process to generate dsRNA to enhance siRNA biogenesis [24, 30, 31]. Therefore, in quinoa, the dominant nucleotide size of 24nt, which are known as hc-siRNAs and had the 5’A bias, were likely generated by association of DCL3, AGO4/6/9, and RDR2. Also, in our analysis presence of other endogenous siRNAs with different sizes (21-27 nt) could be attributed to the function of RDR2. The RDR2 utilizes the stem-loop precursors as the template for production of secondary siRNAs; these precursors are the origin of mature miRNAs [30].

Ten out of eleven known miRNA families identified in this study were also reported at least once elsewhere from *C. quinoa* [4, 99]; miR395 was reported only from this study. This indicated conserved structure of the detected miRNAs between this study and the others, and reliability of our *in silico* miRNA analyses. Identification of known and novel miRNAs in quinoa [4, 99] and prediction of their target genes has already been reported [99]; however, screening of DEmiRNAs in quinoa between CMV- and mock-inoculated samples had not yet been investigated until this study. DE analysis detected a total of 13 DEmiRNAs from quinoa varieties, which overall targeted 126 candidate genes (Table 3, Supplementary Table S6). As miRNA targets 3’ UTR of the target genes for degradation by RISC [100], there are several lines of evidence confirming that there is a strong negative correlation between miRNA expression and regulation of their targeted mRNA [47, 101-103]. In ‘Jessie’, downregulation of novel miRNA “NW_018743850.1_23708” derepressed expression of ABA receptor genes, which are known to modulate ABA levels in plants [104]. Increased ABA may result in diverse defense responses: 1) elevated callose deposition in plasmodesmata to confine and eliminate the virus, and/or 2) a viral strategy to inhibit host systemic resistance via disruption of SA/JA signaling pathway. The role of miR399 in relationship with lipid metabolism have been reported [105]. In ‘Jessie’, suppression of miR399 and miR166, and induction of novel miRNA “NW_018744092.1_28071”, might lead to upregulation of target genes involved in lipid biosynthesis, and one may infer that the virus regulated three miRNAs to modulate host lipid biosynthesis for its own multiplication. Another role of miR399 is to regulate ubiquitin-conjugating E2 enzyme that target proteins for degradation [106]. In ‘Jessie’, downregulation of miR399 may induce: 1) expression of ubiquitin enzyme that increases degradation of viral proteins, and 2) endocytosis, which is a host mechanism to remove ligands, nutrients, lipids and proteins from the cell.

The complexity in siRNA classification during host-virus interaction at genome-wide levels needs development of particular analysis. Using ShortStack facilitated identification of siRNAs based upon 1) software development according to plant genomes, and 2) prediction of *de novo* regions over the genome with accumulated siRNAs named “clusters”. In the *Arabidopsis*-root knot nematode interaction, abundance of 23 and 24 siRNAs is higher in galled samples than in controls and was attributed as a gall characteristic [29, 34]. In this study, however, significant higher accumulation of 24nt siRNA clusters among varieties (‘Red Head’, bitter variety had more siRNA than other two varieties), but not among treatments and harvest times, may suggest that varied abundance of siRNA clusters in quinoa was probably a variety-specific incidence. Because hc-siRNA were the most abundant cleaved size among quinoa siRNA clusters and are associated with silencing of gene and intergenic repeats through DNA methylation and histone modification, differential expression analysis focused only on hc-siRNAs. Similar to miRNAs that have negative correlation with expression of their targets, siRNA expression also influences their targets inversely [29, 107]. Thus, differentially expressed clusters of hc-siRNAs in quinoa either in overall or variety-specific modes impact their putative genomic regions including genes, transcription factor binding sites, and transposable elements in a negative manner. Based on target analyses of differentially expressed hc-siRNA in quinoa, intergenic areas and repetitive elements were more regulated than coding sequences; this is consistent with the fact that hc-siRNA mostly target promoter regions [29]. The differentially expressed hc-siRNAs identified in this study may be good resistance candidates during virus infection, however, further functional studies should be conducted to verify their expression via RNA-directed DNA methylation, and their potential role in quinoa-virus interaction.

During CMV infection in *Arabidopsis*, DCL4 produced 21nt vsiRNA that has been implicated in viral gene silencing. DCL4 mutants had a higher abundance of 22nt vsiRNA, indicating recruitment of DCL2 to compensate DCL4 impairment [25, 28]. Both DCL4 and DCL2 may also be involved in vsiRNA biogenesis in quinoa since 21 and 22 nt are the predominant sizes (Figure 4; Supplementary Figure S4). More vsiRNAs mapped to RNA3 than RNA1 or RNA2 (Figure 5), perhaps due to the fact that RNA3 is the most abundant RNA in infected cells. RNA3 also has shorter sequence length and higher replication rate than RNA1 and RNA2 [108, 109]. Since DCL2 and DCL4 preferentially cleave GC-rich template regions [110] and GC content of CMV RNAs is similar (RNA1: 46.5%; RNA2: 45.9%; and RNA3: 46.9%), this eliminated GC content as the reason for uneven distribution of vsiRNAs across CMV genome. In CMV-infected *Arabidopsis* and *Nicotiana*, RDR1 has the primary role in vsiRNA formation with a 5’ selection bias for three viral genomic RNAs. In RDR1 mutants there was an increased production of vsiRNAs from the 3’ half of the genomic RNAs, particularly in RNA3, which appeared to be RDR6-dependent [109, 111]. Thus, in our study, variability of 5’ bases (Figure 4B) and higher number of vsiRNAs mapped to RNA3 (Figure 5) may have been due to reduced activity of RDR1 and co-expression of RDR6 during vsiRNA formation.

## 5. Conclusion

Altogether, we concluded that CMV infection of quinoa varieties with different saponin profiles resulted in variable virus-induced local or systemic symptoms. Suppression of TSARL1 gene in the bitter variety (high saponin level) was concurrent with systemic movement of the virus in the leaves. Integration of high-throughput transcriptome and sRNASeq of quinoa varieties with different saponin profiles during time-course CMV infection provided the knowledge on 1) general and individual variety responses such as perturbation of translation, lipid and nitrogen metabolism as well as plant-pathogen interaction, hormonal signaling, DNA and mismatch repair processes, and 2) differentially expressed miRNAs and siRNAs and their putative targets. Our findings will enhance knowledge on 1) genes and biological pathways involved in tolerance of quinoa varieties varying in saponin content to viral infection, and 2) molecular characteristics of regulatory miRNAs, endogenous and exogenous siRNAs. However, functional studies should be conducted to confirm the impact of regulatory and endogenous sRNA in quinoa as well as their role in quinoa-virus interaction. Since, the role of saponin in quinoa tolerance to CMV was inconclusive, further research with emphasis on silencing of TSARL1 gene could be of interest to characterize the potential defensive role of saponin in virus infection. Taken together, information provided in this study will be helpful in development of resistant quinoa varieties against virus infection through molecular breeding.

## Supplementary Materials

**Figure S1.** Principal Component Analysis (PCA) based on gene expression of quinoa varieties during time-course cucumber mosaic virus (CMV) infection. The treatments are infected quinoa inoculated with CMV or control as mock-inoculated samples; in varieties, ‘Jessie’ is low saponin (sweet variety), ‘QQ74’ is the medium saponin (sweet variety), and ‘Red Head’ is high saponin (bitter variety); and the time (1 or 4) is the harvesting time post CMV inoculation (dpi). **Figure S2.** TSARL1 expression pattern by qPCR. The “Y” axis shows the log2 fold change of relative expression of TSARL1 gene, and the “X” axis are the varieties abbreviated with initials plus their days-post-inoculation. The red and blue colors indicate high and low expression patterns, respectively. **Figure S3.** siRNA-derived 5’ terminal base abundance. Quinoa varieties (J,Q,R) with two treatments (C,T) at two time periods (days) after inoculation (1,4) are shown in the chart title. Nucleotide size and dominance percentage of quinoa reads are shown in x and y axis, respectively. **Figure S4.** vsiRNA-derived 5’ terminal base abundance. Quinoa varieties (J,Q,R) inoculated with CMV (T) at two time periods (days) after inoculation (1,4) are shown in the chart title. Nucleotide size and dominance percentage of viral reads are shown in x and y axis, respectively. **Table S1.** Primers used in validation of DEGs, known and novel miRNAs. **Table S2.** Statistics of RNASeq and small RNASeq data. Number of raw, trimmed, and mapped reads per individual library is indicated in respective tab. **Table S3.** Differential expressed genes of RNASeq data. In each tab, “full-3way” indicates full-model analysis, and “full-treatment” indicates main effect of treatment only, upregulated and downregulated indicates comparison of CMV-inoculated and mock-inoculated samples. “Variety-[Jessie, QQ74, Red Head]-2way” means individual variety-specific model while considering time and treatment effect, whereas “Variety-[Jessie, QQ74, Red Head]-treatment means individual variety-specific model by considering main effect of treatment only. upregulated and downregulated indicates comparison of CMV-inoculated and mock-inoculated samples. **Table S4.** Gene annotation and biological function of RNASeq DEGs obtained through GO term and KEGG pathways analyses. For tab descriptions refer to Table S3. **Table S5.** Sequences and statistics of predicted known and novel (mature, star, and precursor) miRNA utilizing miRDeep2. **Table S6.** Biological functions of the genes that are targeted by differential expressed miRNAs (DEmiRNA). Since characterization of DEmiRNA was followed as the DEGs from RNASeq, thus, for description of the tabs refer to Table S3. Note that in this file, up/downregulation is pertained to expression pattern of DEmiRNAs. **Table S7.** ShortStack-generated statistics about siRNA clusters. Genome coordination, length, number of reads mapped in million (RPM), unique reads, cleaved sizes, and abundance of cleaved nucleotide of the clusters are shown. **Table S8.** Information of differential expressed siRNA cluster based on full or variety-specific models. **Table S9.** Predicted targets (gene function, biological pathway, transcription factor binding site [TFBS], transposable element [TE]) of differential expressed siRNA (DEsiRNA). For tab descriptions refer to Table S3. Note that in this file, up/downregulation pattern is pertained to expression pattern of DEsiRNAs.

## Author Contributions

Conceptualization, Nourolah Soltani, Margaret Staton and Kimberly Gwinn; Data curation, Nourolah Soltani; Formal analysis, Nourolah Soltani and Margaret Staton; Funding acquisition, Kimberly Gwinn; Investigation, Nourolah Soltani; Methodology, Nourolah Soltani, Margaret Staton and Kimberly Gwinn; Project administration, Kimberly Gwinn; Resources, Nourolah Soltani, Margaret Staton and Kimberly Gwinn; Software, Nourolah Soltani; Supervision, Margaret Staton and Kimberly Gwinn; Validation, Nourolah Soltani, Margaret Staton and Kimberly Gwinn; Visualization, Nourolah Soltani; Writing – original draft, Nourolah Soltani; Writing – review & editing, Margaret Staton and Kimberly Gwinn.. All authors have read and agreed to the published version of the manuscript.

## Funding

This work was supported in part by the USDA National Institute of Food and Agriculture, Hatch project TEN0048 (K.D.G.). Research on small RNA was also supported by a grant to N.S. and K.D.G. from the University of Tennessee Graduate School.

## Conflicts of Interest

The authors declare no conflict of interest

## Acknowledgments

Authors thank Jason Abbott and Jim Hermann for the generous gift of seed of quinoa variety ‘Jessie’ and providing information on the genetics of the variety. We also thank the U.S. National Plant Germplasm System – North Central Plant Introduction Station for seed of quinoa ‘QQ74’. In addition, we thank Frank Morton for information on the genetics of ‘Red Head’. This work is part of the doctoral dissertation of Nourolah Soltani, “Genome-enabled analysis of *Quercus rubra*-ozone and *Chenopodium quinoa*-cucumber mosaic virus interactions” (University of Tennessee).

## Notes

### Competing Interest Statement

The authors have declared no competing interest.

