## Supplementary material for "Integration of Transcriptome and Small RNA Sequencing to Decipher Molecular Interaction of *Chenopodium quinoa* Varieties with Cucumber Mosaic Virus": Supplementary Figure S3 - siRNA 5' terminal base abundance.pptx

### Slide 1
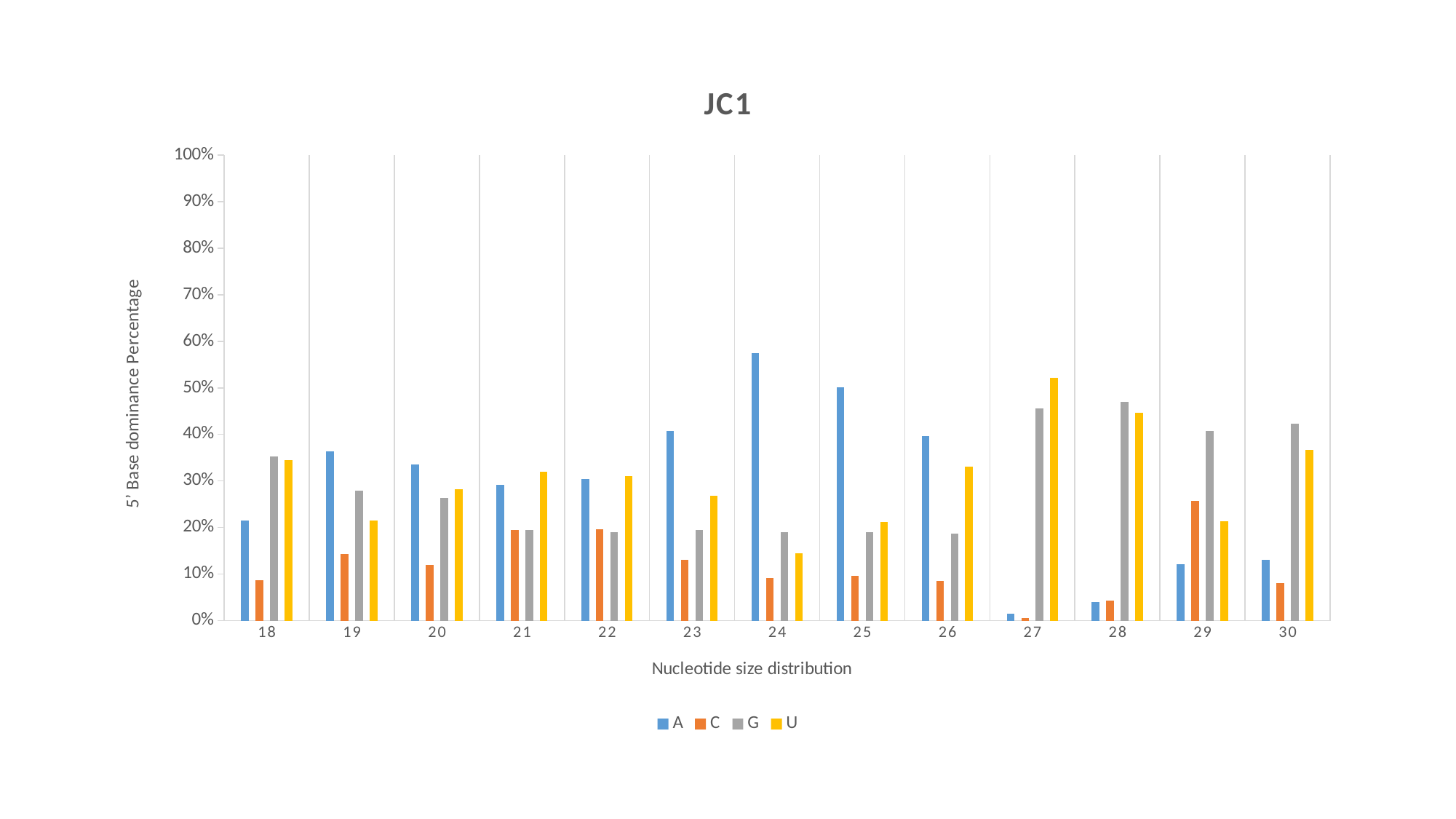

#### Chart: JC1
| Category | A | C | G | U |
|---|---|---|---|---|
| 18 | 0.215283 | 0.086747 | 0.352547 | 0.345423 |
| 19 | 0.36348 | 0.14261666666666667 | 0.27884 | 0.21505666666666667 |
| 20 | 0.33503333333333335 | 0.11947999999999999 | 0.26378666666666667 | 0.2817 |
| 21 | 0.2910933333333333 | 0.19419666666666668 | 0.19529333333333332 | 0.3194166666666667 |
| 22 | 0.30462000000000006 | 0.19597333333333333 | 0.18931666666666666 | 0.31009 |
| 23 | 0.40707999999999994 | 0.13023666666666667 | 0.19478666666666666 | 0.2679 |
| 24 | 0.5745833333333333 | 0.09116999999999999 | 0.18952 | 0.14472666666666667 |
| 25 | 0.5017266666666668 | 0.09548 | 0.19015 | 0.21264333333333332 |
| 26 | 0.39697333333333334 | 0.08441333333333333 | 0.18755999999999998 | 0.33105999999999997 |
| 27 | 0.015543333333333334 | 0.006273333333333333 | 0.45587333333333335 | 0.52231 |
| 28 | 0.03937333333333334 | 0.04365 | 0.47024333333333335 | 0.44673999999999997 |
| 29 | 0.12150000000000001 | 0.25798333333333334 | 0.4073833333333334 | 0.21312999999999996 |
| 30 | 0.13009666666666667 | 0.08038333333333333 | 0.42236 | 0.36716 |

### Slide 2
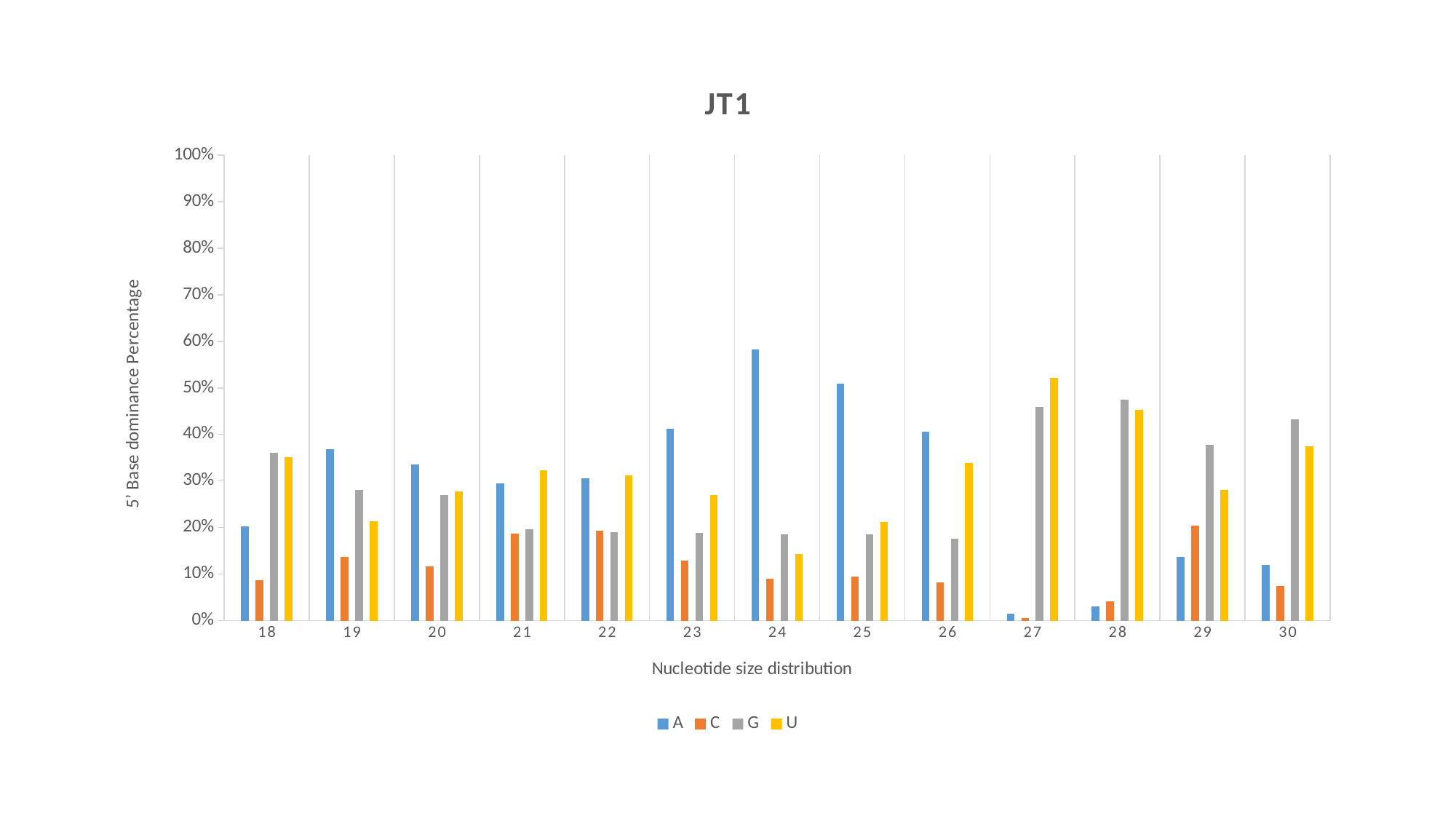

#### Chart: JT1
| Category | A | C | G | U |
|---|---|---|---|---|
| 18 | 0.20253999999999997 | 0.08642666666666667 | 0.3601666666666667 | 0.35087 |
| 19 | 0.36750666666666665 | 0.13712333333333335 | 0.28147666666666665 | 0.21389666666666665 |
| 20 | 0.33515999999999996 | 0.11620000000000001 | 0.27047 | 0.27817000000000003 |
| 21 | 0.29436666666666667 | 0.1867466666666667 | 0.19656333333333334 | 0.32232 |
| 22 | 0.3056933333333334 | 0.19276333333333331 | 0.19017666666666666 | 0.3113666666666666 |
| 23 | 0.41243 | 0.12888999999999998 | 0.18893333333333331 | 0.26974333333333333 |
| 24 | 0.5821 | 0.08995666666666667 | 0.18541333333333335 | 0.14253333333333332 |
| 25 | 0.50876 | 0.09494999999999999 | 0.18457333333333334 | 0.21171333333333334 |
| 26 | 0.4050766666666667 | 0.08129333333333333 | 0.17572333333333334 | 0.33790666666666663 |
| 27 | 0.014039999999999999 | 0.006176666666666667 | 0.45821666666666666 | 0.5215666666666667 |
| 28 | 0.031023333333333337 | 0.041166666666666664 | 0.47451 | 0.4532966666666667 |
| 29 | 0.13752333333333333 | 0.20374666666666666 | 0.37778 | 0.2809466666666667 |
| 30 | 0.11965333333333333 | 0.07451 | 0.43198000000000003 | 0.3738566666666667 |

### Slide 3
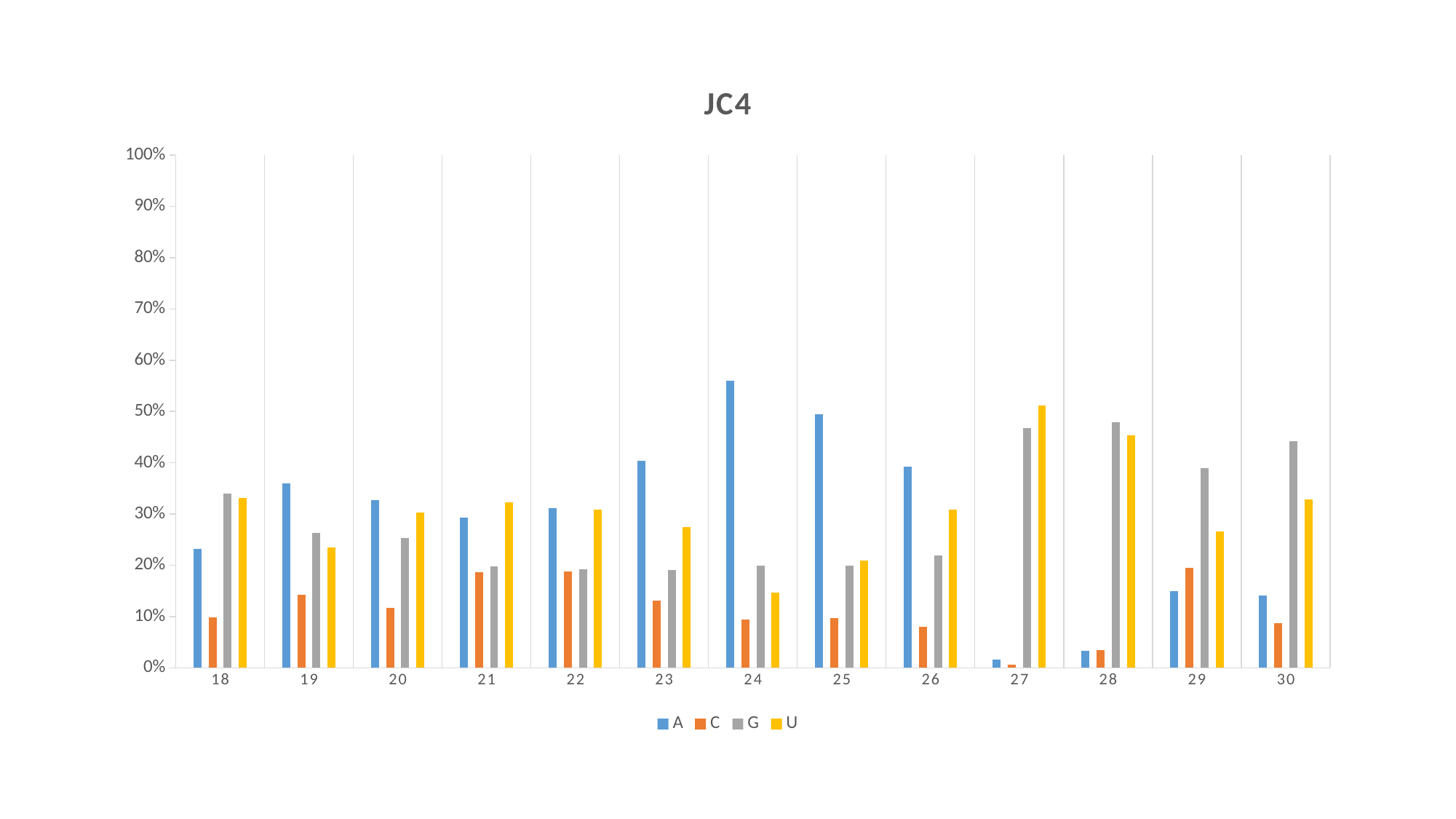

#### Chart: JC4
| Category | A | C | G | U |
|---|---|---|---|---|
| 18 | 0.23140666666666665 | 0.09806999999999999 | 0.3394 | 0.33112 |
| 19 | 0.35985333333333336 | 0.14233 | 0.26305333333333336 | 0.23476666666666668 |
| 20 | 0.3268333333333333 | 0.11724666666666667 | 0.2533666666666667 | 0.3025533333333334 |
| 21 | 0.2936033333333334 | 0.18652000000000002 | 0.19752999999999998 | 0.32234666666666667 |
| 22 | 0.31137 | 0.18761000000000003 | 0.19259 | 0.30843333333333334 |
| 23 | 0.40365 | 0.13047333333333333 | 0.19104666666666667 | 0.27482999999999996 |
| 24 | 0.5604966666666666 | 0.09423 | 0.19884 | 0.14643666666666666 |
| 25 | 0.49525 | 0.09669666666666665 | 0.19876333333333332 | 0.20928999999999998 |
| 26 | 0.39308000000000004 | 0.07977333333333335 | 0.21905333333333332 | 0.30809333333333333 |
| 27 | 0.015556666666666665 | 0.005749999999999999 | 0.4674466666666666 | 0.5112466666666666 |
| 28 | 0.03322 | 0.034416666666666665 | 0.4787266666666667 | 0.4536366666666667 |
| 29 | 0.14995 | 0.19516999999999998 | 0.38933999999999996 | 0.2655433333333333 |
| 30 | 0.14156000000000002 | 0.08764333333333334 | 0.4415233333333333 | 0.32926999999999995 |

### Slide 4
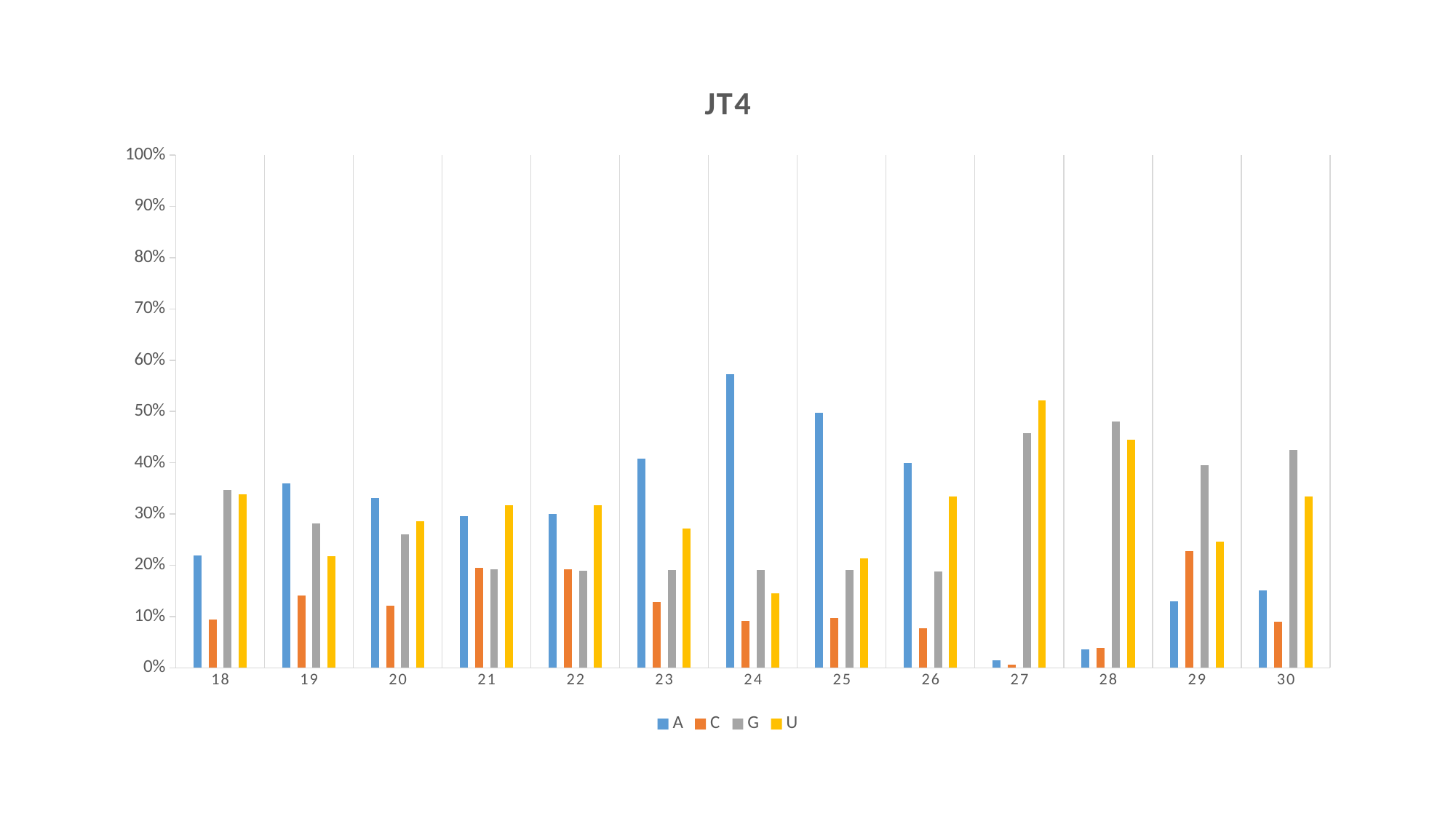

#### Chart: JT4
| Category | A | C | G | U |
|---|---|---|---|---|
| 18 | 0.21924 | 0.09437333333333335 | 0.34741 | 0.33897333333333335 |
| 19 | 0.35978333333333334 | 0.14142666666666667 | 0.2816533333333333 | 0.21713000000000002 |
| 20 | 0.33193333333333336 | 0.12123333333333335 | 0.26035333333333327 | 0.28648333333333337 |
| 21 | 0.29531666666666667 | 0.19442333333333336 | 0.19245666666666664 | 0.31780333333333327 |
| 22 | 0.30074333333333336 | 0.19161333333333333 | 0.19005333333333332 | 0.31759 |
| 23 | 0.40873 | 0.12856666666666666 | 0.1909633333333333 | 0.27174000000000004 |
| 24 | 0.57255 | 0.09116666666666666 | 0.19036333333333333 | 0.14592000000000002 |
| 25 | 0.4982066666666667 | 0.09735333333333333 | 0.19036 | 0.2140766666666667 |
| 26 | 0.3994866666666666 | 0.07776666666666666 | 0.18835000000000002 | 0.3343966666666667 |
| 27 | 0.01462 | 0.006430000000000001 | 0.4578 | 0.5211466666666666 |
| 28 | 0.03601666666666667 | 0.038766666666666665 | 0.48095666666666664 | 0.44426000000000004 |
| 29 | 0.13003333333333333 | 0.22762333333333332 | 0.3955933333333333 | 0.24675000000000002 |
| 30 | 0.15159 | 0.09000666666666667 | 0.4245666666666667 | 0.3338366666666667 |

### Slide 5
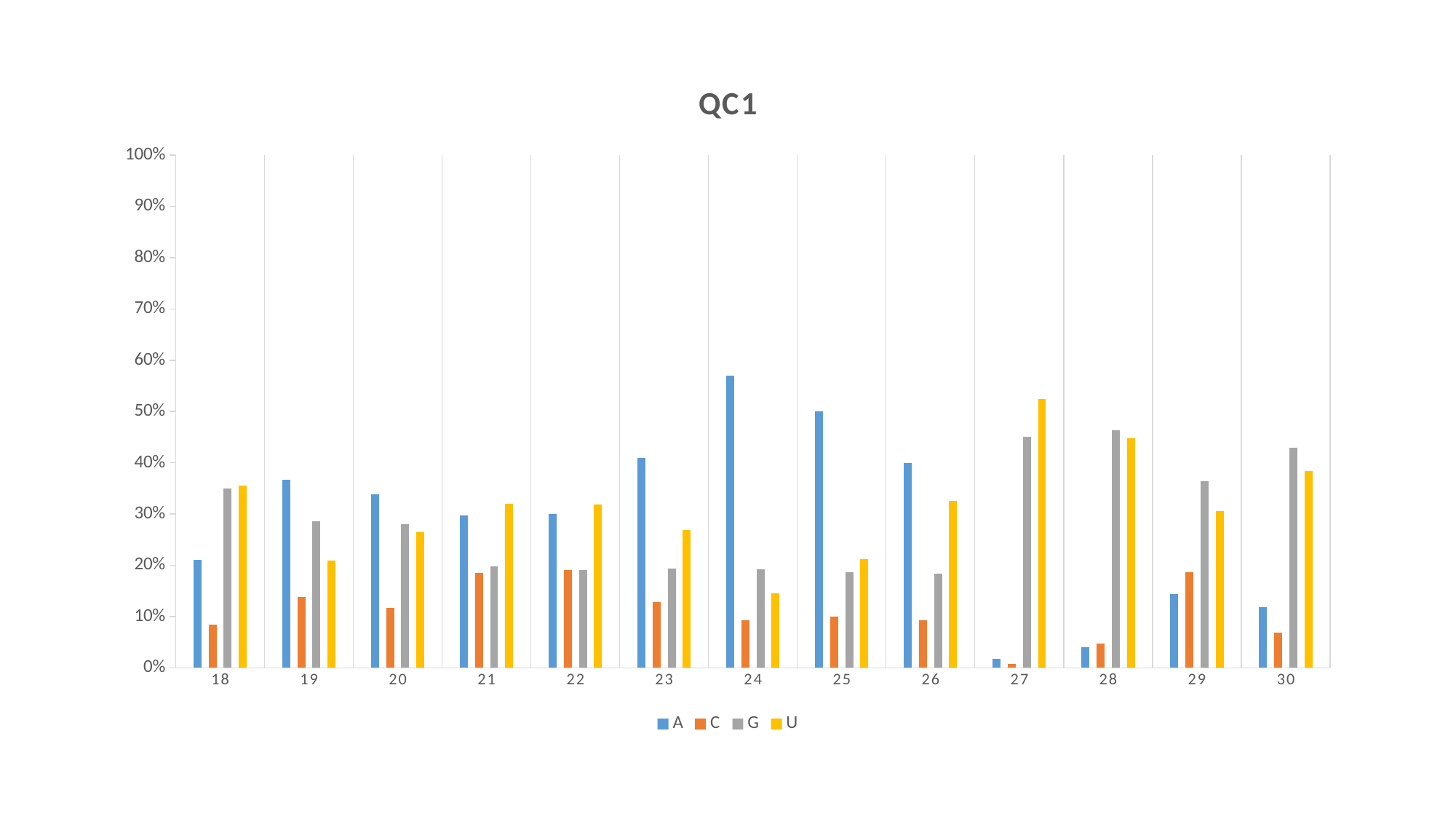

#### Chart: QC1
| Category | A | C | G | U |
|---|---|---|---|---|
| 18 | 0.21063333333333334 | 0.08416333333333333 | 0.3499366666666666 | 0.3552633333333333 |
| 19 | 0.3663 | 0.13763 | 0.28633333333333333 | 0.20973333333333333 |
| 20 | 0.33843 | 0.11649666666666665 | 0.27988 | 0.2651966666666667 |
| 21 | 0.2976333333333333 | 0.1847633333333333 | 0.19754666666666665 | 0.3200533333333333 |
| 22 | 0.3006466666666667 | 0.19022666666666668 | 0.19088333333333332 | 0.3182433333333334 |
| 23 | 0.40972666666666663 | 0.12800666666666669 | 0.19360333333333335 | 0.26866666666666666 |
| 24 | 0.57006 | 0.09276000000000001 | 0.19203333333333336 | 0.14514666666666667 |
| 25 | 0.49998666666666663 | 0.10015 | 0.18722333333333332 | 0.21264000000000002 |
| 26 | 0.3992 | 0.09262 | 0.18314 | 0.32504 |
| 27 | 0.01758 | 0.007423333333333333 | 0.45038666666666666 | 0.5246066666666668 |
| 28 | 0.04043333333333333 | 0.047966666666666664 | 0.46334666666666663 | 0.44826000000000005 |
| 29 | 0.14347666666666667 | 0.1859833333333333 | 0.36416 | 0.3063766666666667 |
| 30 | 0.11809333333333333 | 0.06819333333333333 | 0.4294766666666667 | 0.3842366666666667 |

### Slide 6
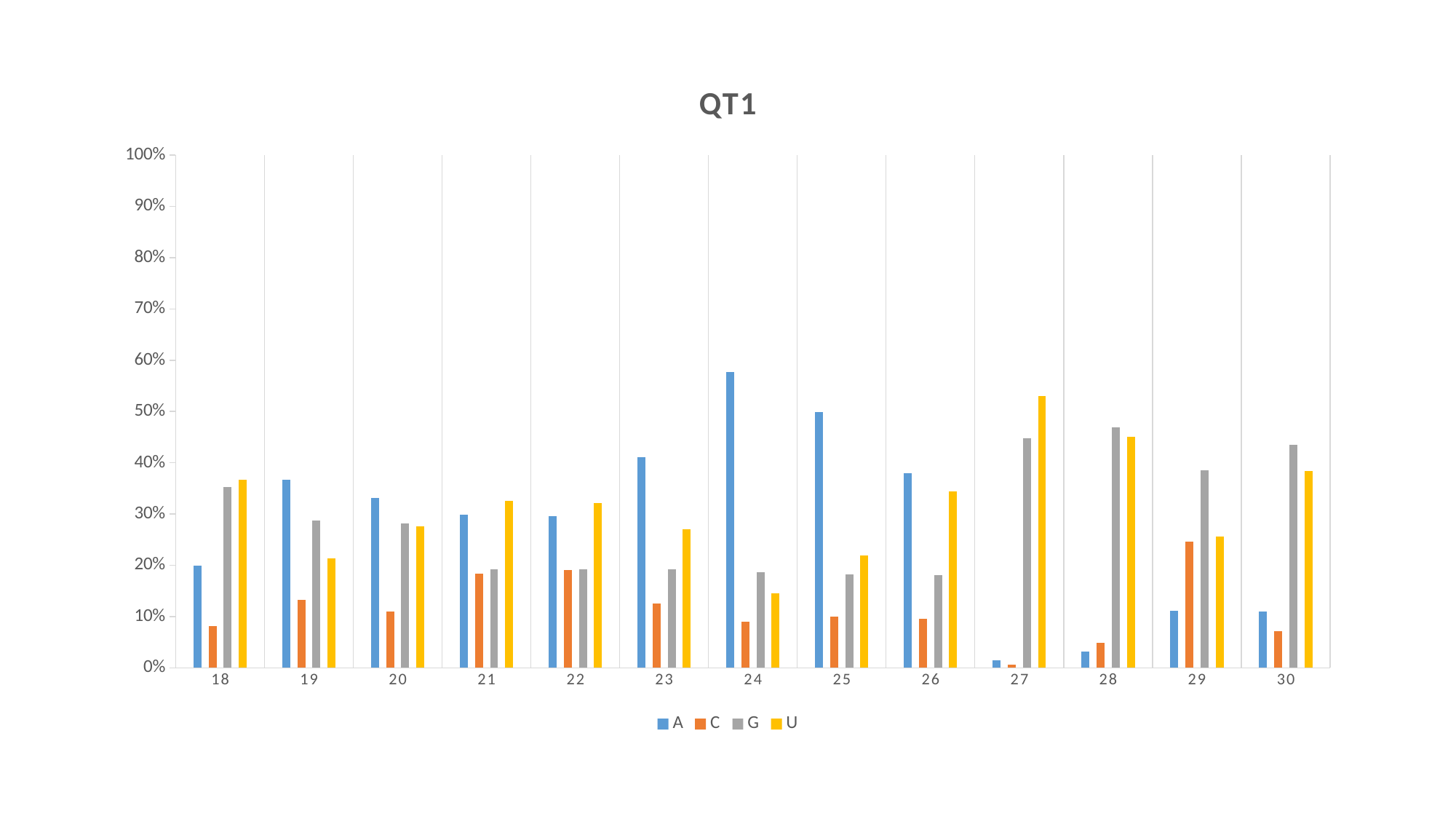

#### Chart: QT1
| Category | A | C | G | U |
|---|---|---|---|---|
| 18 | 0.19882666666666668 | 0.08185 | 0.3524433333333333 | 0.36688000000000004 |
| 19 | 0.36677 | 0.1321 | 0.28761333333333333 | 0.21351333333333333 |
| 20 | 0.33189 | 0.10937333333333332 | 0.2820933333333333 | 0.27664000000000005 |
| 21 | 0.29906666666666665 | 0.18337666666666666 | 0.19205333333333333 | 0.32550333333333337 |
| 22 | 0.29624999999999996 | 0.19019333333333333 | 0.19188666666666668 | 0.3216666666666667 |
| 23 | 0.41118666666666664 | 0.12506666666666666 | 0.1929166666666667 | 0.27082333333333336 |
| 24 | 0.5774 | 0.08961999999999999 | 0.18717000000000003 | 0.14581333333333335 |
| 25 | 0.4988966666666667 | 0.09936666666666666 | 0.18225333333333335 | 0.21948333333333334 |
| 26 | 0.37914000000000003 | 0.09516333333333334 | 0.18083666666666665 | 0.34486333333333336 |
| 27 | 0.014536666666666668 | 0.006493333333333334 | 0.44817333333333337 | 0.5308066666666668 |
| 28 | 0.03173666666666667 | 0.048850000000000005 | 0.4687966666666667 | 0.45062 |
| 29 | 0.11155333333333334 | 0.24689999999999998 | 0.38555333333333336 | 0.25599 |
| 30 | 0.11036666666666667 | 0.07163333333333334 | 0.43477666666666664 | 0.38322666666666666 |

### Slide 7
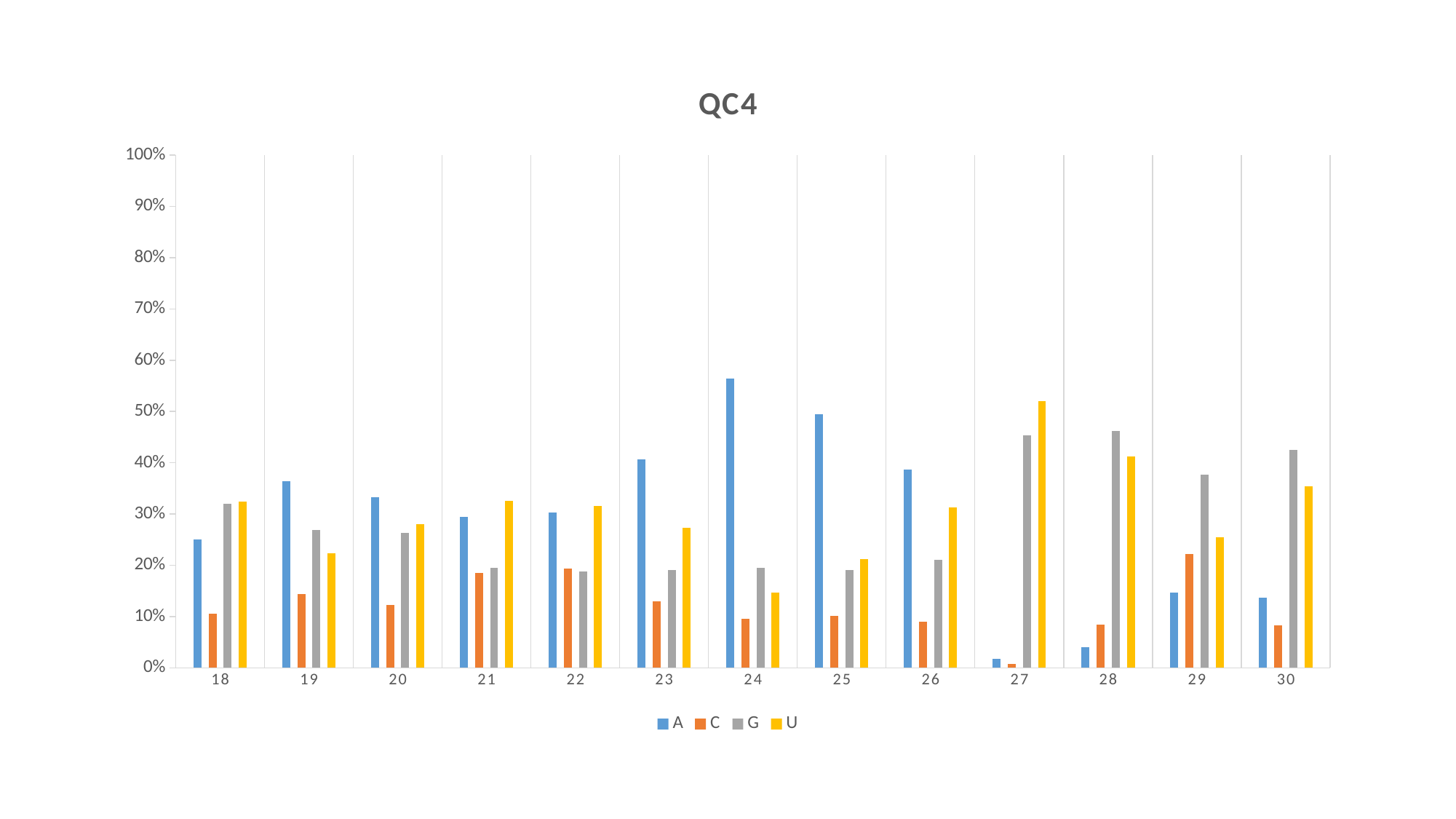

#### Chart: QC4
| Category | A | C | G | U |
|---|---|---|---|---|
| 18 | 0.25044 | 0.10541333333333334 | 0.31965333333333334 | 0.32449 |
| 19 | 0.36365333333333333 | 0.14398333333333332 | 0.26881 | 0.22355666666666665 |
| 20 | 0.33226 | 0.12318 | 0.2638933333333333 | 0.2806633333333333 |
| 21 | 0.29440666666666665 | 0.1856266666666667 | 0.19447666666666666 | 0.32548666666666665 |
| 22 | 0.3028566666666667 | 0.19299999999999998 | 0.18798666666666666 | 0.31615333333333334 |
| 23 | 0.4061966666666666 | 0.13034666666666667 | 0.19042666666666666 | 0.2730333333333333 |
| 24 | 0.5641766666666667 | 0.09512333333333334 | 0.1944233333333333 | 0.1462766666666667 |
| 25 | 0.49491666666666667 | 0.10167666666666665 | 0.19142333333333336 | 0.21198333333333333 |
| 26 | 0.38714333333333334 | 0.09009 | 0.21025333333333332 | 0.31251 |
| 27 | 0.017806666666666665 | 0.007673333333333333 | 0.45402666666666663 | 0.5204933333333334 |
| 28 | 0.04078 | 0.085 | 0.4624066666666667 | 0.41181 |
| 29 | 0.14649333333333334 | 0.22221000000000002 | 0.3765833333333333 | 0.25471333333333335 |
| 30 | 0.13734333333333335 | 0.08337666666666665 | 0.4255533333333334 | 0.3537266666666667 |

### Slide 8
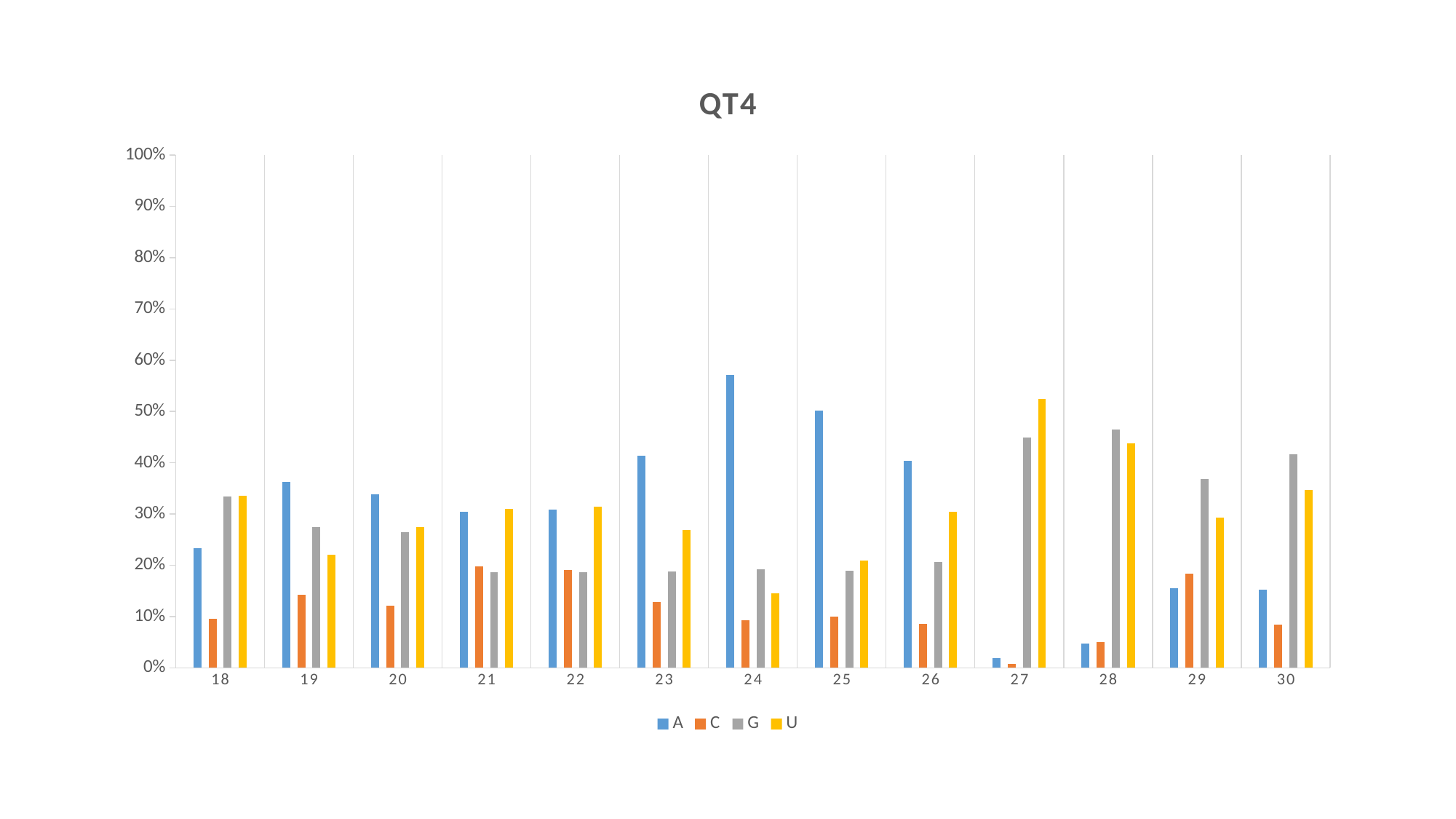

#### Chart: QT4
| Category | A | C | G | U |
|---|---|---|---|---|
| 18 | 0.23406666666666665 | 0.09538666666666668 | 0.33472999999999997 | 0.3358133333333333 |
| 19 | 0.36203 | 0.14222333333333334 | 0.2750633333333333 | 0.22068 |
| 20 | 0.33896666666666664 | 0.12145333333333334 | 0.2651233333333333 | 0.2744566666666666 |
| 21 | 0.30487333333333333 | 0.19816333333333333 | 0.18666000000000002 | 0.3103033333333333 |
| 22 | 0.30929333333333336 | 0.19054666666666664 | 0.18633 | 0.31383 |
| 23 | 0.41417 | 0.12861666666666668 | 0.18782666666666667 | 0.26938666666666666 |
| 24 | 0.5710633333333334 | 0.09229666666666665 | 0.19185 | 0.14479 |
| 25 | 0.5018966666666667 | 0.09947 | 0.18954666666666667 | 0.20908000000000002 |
| 26 | 0.40326 | 0.08538333333333332 | 0.2068566666666667 | 0.3045 |
| 27 | 0.01861 | 0.00751 | 0.4486033333333333 | 0.5252766666666667 |
| 28 | 0.04706 | 0.04968 | 0.465 | 0.4382633333333333 |
| 29 | 0.15553666666666668 | 0.18394 | 0.3676733333333333 | 0.29285666666666665 |
| 30 | 0.15196666666666667 | 0.08486 | 0.41673 | 0.34644 |

### Slide 9
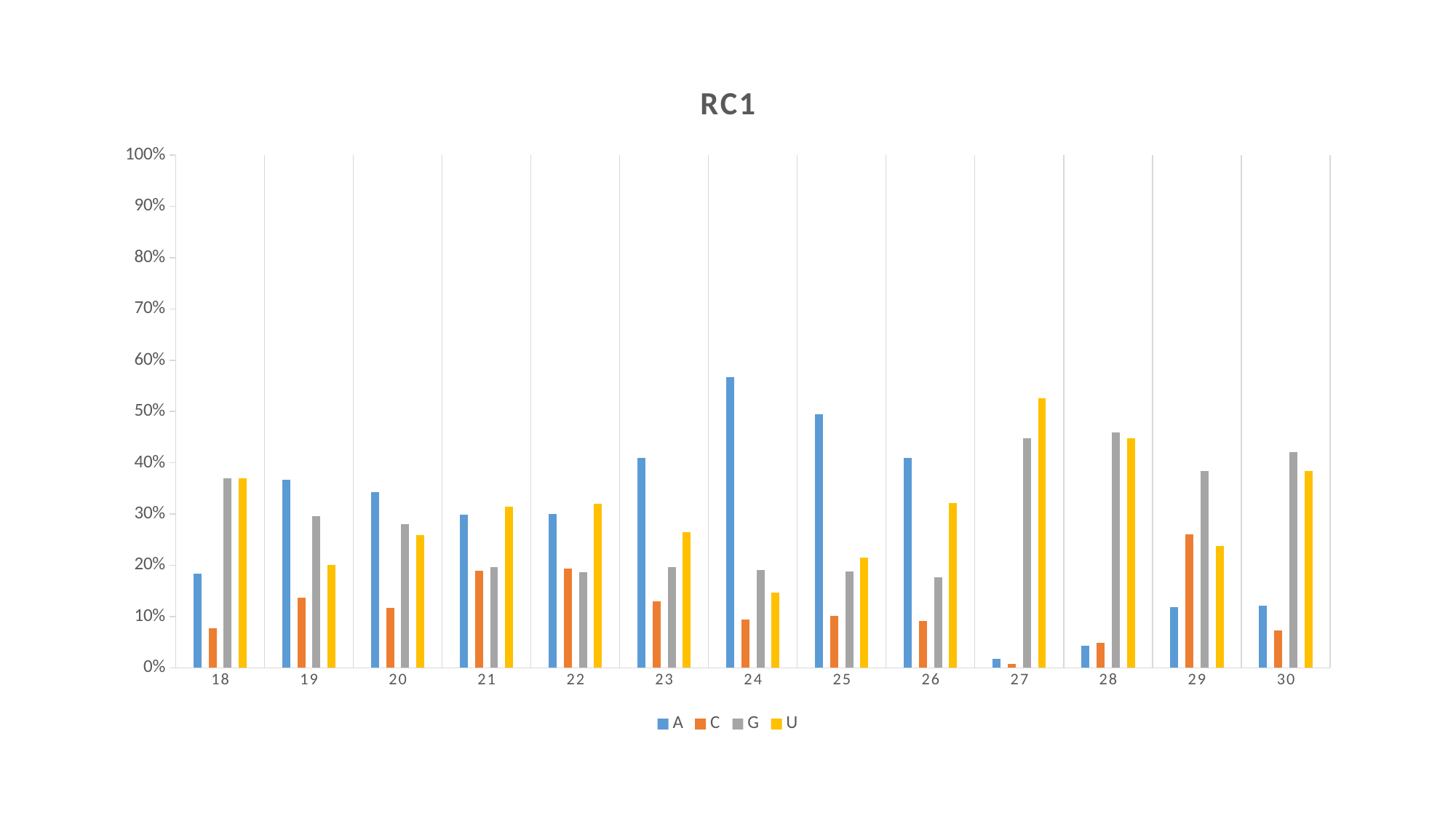

#### Chart: RC1
| Category | A | C | G | U |
|---|---|---|---|---|
| 18 | 0.18350666666666668 | 0.07729333333333333 | 0.36927666666666664 | 0.3699266666666667 |
| 19 | 0.36744 | 0.13618666666666668 | 0.29522000000000004 | 0.20114666666666667 |
| 20 | 0.34324333333333334 | 0.11733333333333333 | 0.27991666666666665 | 0.25950333333333336 |
| 21 | 0.2991833333333333 | 0.18958 | 0.19643333333333332 | 0.3148033333333333 |
| 22 | 0.3004 | 0.19366666666666668 | 0.18642 | 0.31951 |
| 23 | 0.40981666666666666 | 0.12926 | 0.19666333333333333 | 0.2642566666666667 |
| 24 | 0.5676066666666667 | 0.09422333333333333 | 0.19094 | 0.14723 |
| 25 | 0.49538 | 0.10133333333333333 | 0.18804333333333334 | 0.21524333333333331 |
| 26 | 0.40996666666666665 | 0.0909 | 0.17726666666666668 | 0.3218666666666667 |
| 27 | 0.01803 | 0.007936666666666667 | 0.44851 | 0.5255299999999999 |
| 28 | 0.04354666666666667 | 0.049229999999999996 | 0.45927 | 0.44794666666666666 |
| 29 | 0.11795333333333334 | 0.26048333333333334 | 0.3838433333333333 | 0.23772000000000001 |
| 30 | 0.12178666666666667 | 0.07243 | 0.42135999999999996 | 0.38442333333333334 |

### Slide 10
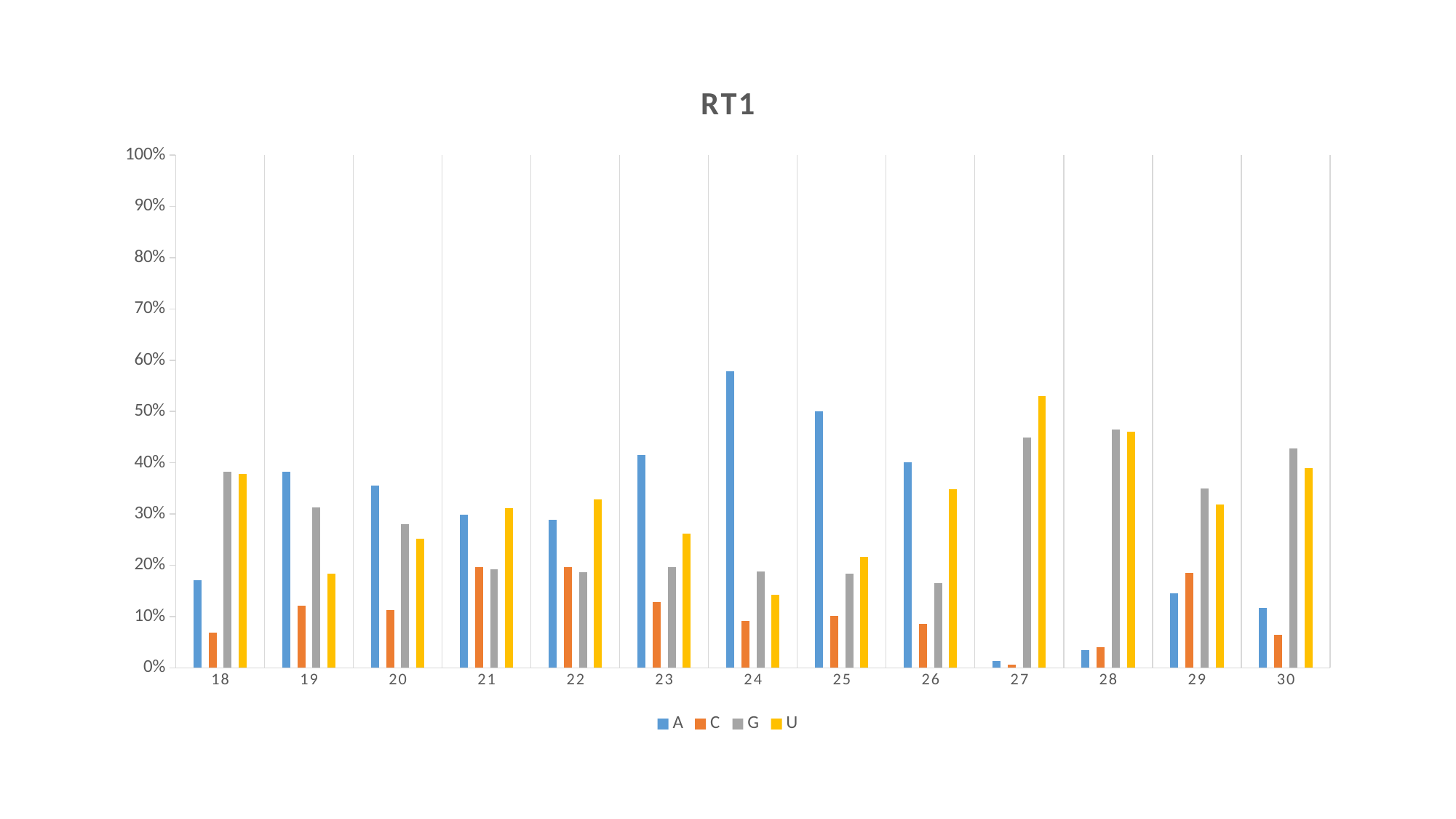

#### Chart: RT1
| Category | A | C | G | U |
|---|---|---|---|---|
| 18 | 0.17078666666666664 | 0.06880000000000001 | 0.3818533333333334 | 0.37856 |
| 19 | 0.38304333333333335 | 0.12072 | 0.31231666666666663 | 0.18392666666666668 |
| 20 | 0.3553566666666666 | 0.11344333333333334 | 0.27960999999999997 | 0.25159333333333334 |
| 21 | 0.29904 | 0.1966 | 0.19244666666666665 | 0.31190999999999997 |
| 22 | 0.28830333333333336 | 0.19580333333333333 | 0.18681999999999999 | 0.32907333333333333 |
| 23 | 0.4148266666666667 | 0.12783666666666668 | 0.19585666666666665 | 0.26148 |
| 24 | 0.5781566666666668 | 0.09135 | 0.18794999999999998 | 0.14254 |
| 25 | 0.5000166666666667 | 0.10087 | 0.18330333333333335 | 0.21580999999999997 |
| 26 | 0.40076333333333336 | 0.08605333333333333 | 0.16504333333333335 | 0.3481433333333333 |
| 27 | 0.013806666666666667 | 0.006450000000000001 | 0.4497633333333333 | 0.5299766666666667 |
| 28 | 0.034179999999999995 | 0.04092666666666667 | 0.46451666666666663 | 0.46038 |
| 29 | 0.14595666666666668 | 0.18472 | 0.35051000000000004 | 0.31881333333333334 |
| 30 | 0.11740666666666666 | 0.06476 | 0.42768 | 0.39015 |

### Slide 11
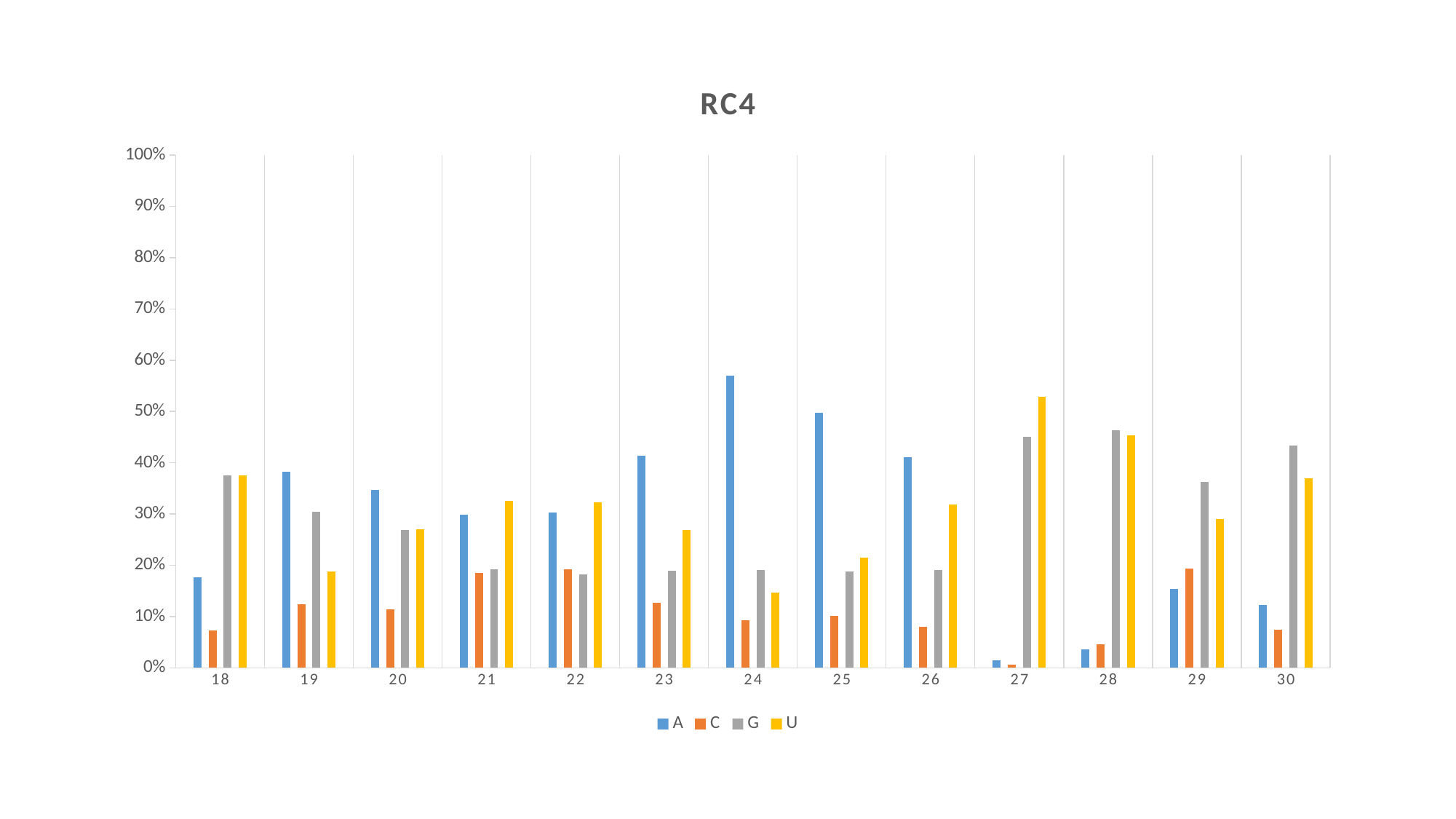

#### Chart: RC4
| Category | A | C | G | U |
|---|---|---|---|---|
| 18 | 0.17635333333333333 | 0.07342333333333334 | 0.3752166666666667 | 0.37501 |
| 19 | 0.38249666666666665 | 0.1245 | 0.30438000000000004 | 0.18862333333333334 |
| 20 | 0.3467766666666667 | 0.11402333333333332 | 0.26848333333333335 | 0.27071666666666666 |
| 21 | 0.29809 | 0.18456666666666668 | 0.19191333333333335 | 0.3254266666666667 |
| 22 | 0.30244000000000004 | 0.19232000000000002 | 0.18220999999999998 | 0.32303333333333334 |
| 23 | 0.4141733333333333 | 0.12756666666666666 | 0.18885 | 0.26940333333333333 |
| 24 | 0.5695100000000001 | 0.09279333333333332 | 0.19064 | 0.14705666666666667 |
| 25 | 0.49704666666666664 | 0.10099999999999999 | 0.18735 | 0.21460333333333334 |
| 26 | 0.4107066666666667 | 0.07978999999999999 | 0.19077666666666668 | 0.31872666666666666 |
| 27 | 0.014536666666666665 | 0.006089999999999999 | 0.4505266666666667 | 0.5288466666666666 |
| 28 | 0.03664666666666666 | 0.04545 | 0.46379333333333334 | 0.45411333333333337 |
| 29 | 0.15338 | 0.19393666666666667 | 0.36194333333333334 | 0.29074 |
| 30 | 0.12280666666666666 | 0.07431333333333334 | 0.43326 | 0.36962 |

### Slide 12
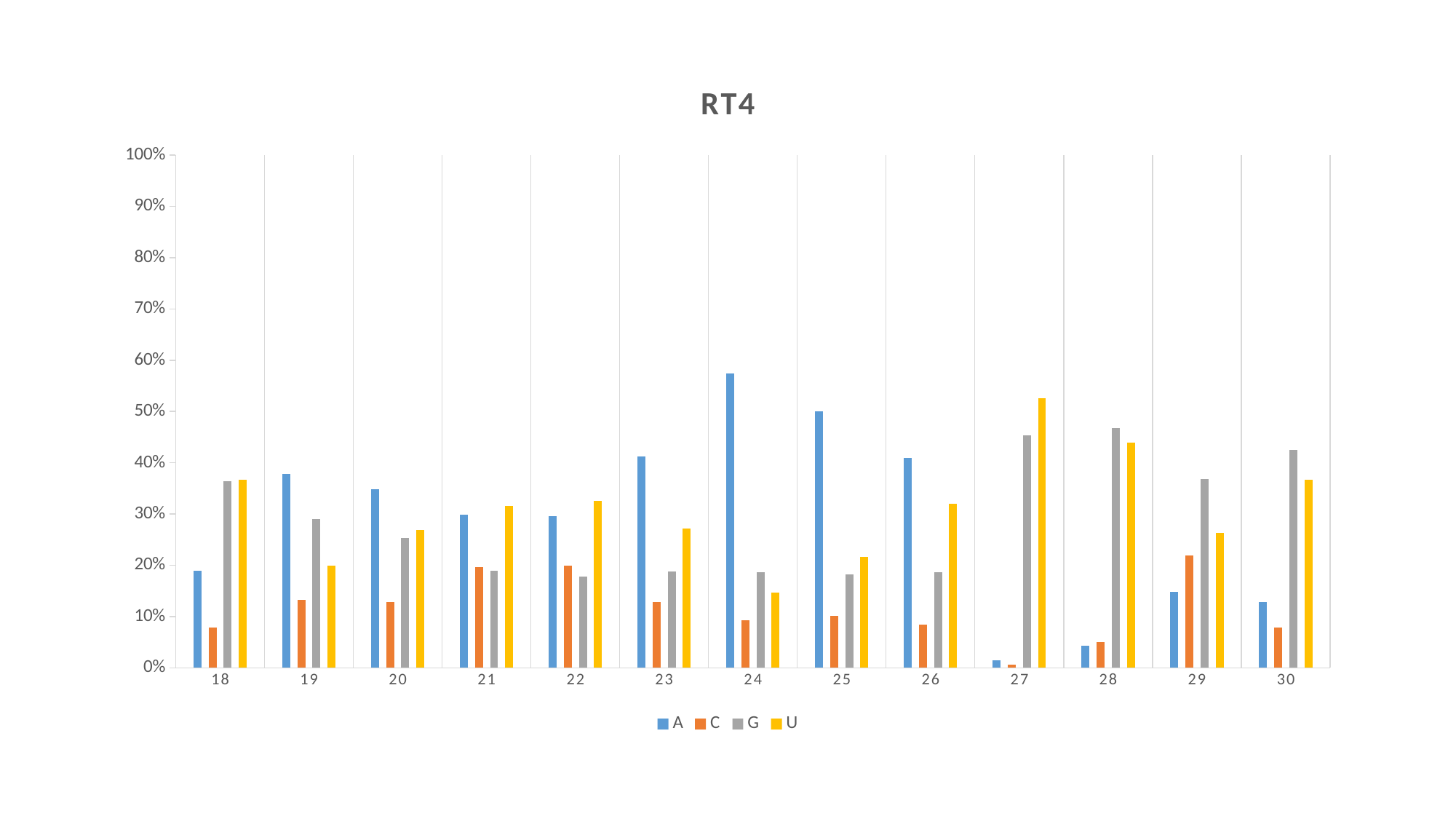

#### Chart: RT4
| Category | A | C | G | U |
|---|---|---|---|---|
| 18 | 0.190056667 | 0.078346667 | 0.364476667 | 0.36712 |
| 19 | 0.37809 | 0.13196 | 0.29038 | 0.19957 |
| 20 | 0.34872 | 0.12861 | 0.253523333 | 0.269143333 |
| 21 | 0.298806667 | 0.196816667 | 0.189006667 | 0.31537 |
| 22 | 0.295606667 | 0.199743333 | 0.178533333 | 0.32612 |
| 23 | 0.4124133333333333 | 0.12868 | 0.18783333333333332 | 0.27107333333333333 |
| 24 | 0.57405 | 0.092353333 | 0.186806667 | 0.14679 |
| 25 | 0.499846667 | 0.10084 | 0.182783333 | 0.216533333 |
| 26 | 0.40953 | 0.083826667 | 0.186356667 | 0.320286667 |
| 27 | 0.014966666666666668 | 0.00632 | 0.45304666666666665 | 0.5256733333333333 |
| 28 | 0.04296 | 0.049826666666666665 | 0.46717999999999993 | 0.4400266666666666 |
| 29 | 0.14871333333333334 | 0.21866333333333332 | 0.3687266666666667 | 0.2638933333333333 |
| 30 | 0.12874333333333335 | 0.07875333333333334 | 0.42573666666666665 | 0.3667666666666667 |
