## Supplementary material for "Integration of Transcriptome and Small RNA Sequencing to Decipher Molecular Interaction of *Chenopodium quinoa* Varieties with Cucumber Mosaic Virus": Supplementary Figure S4 - vsiRNA 5' terminal base abundance.pptx

### Slide 1
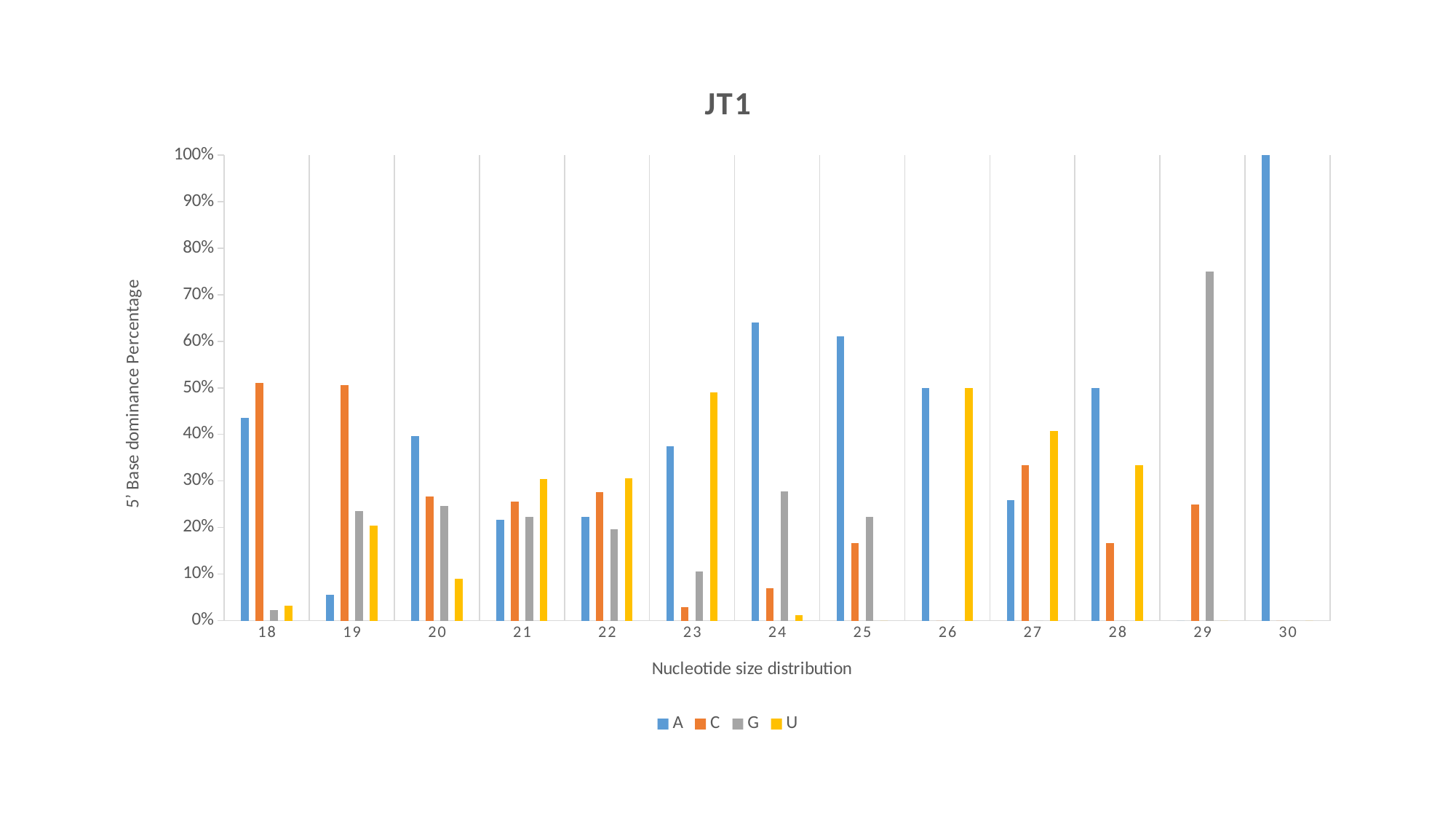

#### Chart: JT1
| Category | A | C | G | U |
|---|---|---|---|---|
| 18 | 0.4349211111111111 | 0.5111111111111111 | 0.022222222222222223 | 0.03174555555555556 |
| 19 | 0.05555555555555555 | 0.5055555555555555 | 0.23518555555555556 | 0.20370333333333335 |
| 20 | 0.3970622222222222 | 0.2667555555555556 | 0.24679777777777775 | 0.08938444444444445 |
| 21 | 0.2164677777777778 | 0.2562488888888889 | 0.22350444444444442 | 0.30378 |
| 22 | 0.22251333333333334 | 0.27538999999999997 | 0.19594111111111112 | 0.30615555555555557 |
| 23 | 0.375 | 0.028703333333333334 | 0.10555555555555556 | 0.4907411111111111 |
| 24 | 0.6407411111111111 | 0.07037 | 0.27777777777777773 | 0.011111111111111112 |
| 25 | 0.61111 | 0.16666666666666666 | 0.22222333333333333 | 0.0 |
| 26 | 0.5 | 0.0 | 0.0 | 0.5 |
| 27 | 0.25925888888888887 | 0.3333333333333333 | 0.0 | 0.4074077777777778 |
| 28 | 0.5 | 0.16666666666666666 | 0.0 | 0.3333333333333333 |
| 29 | 0.0 | 0.25 | 0.75 | 0.0 |
| 30 | 1.0 | 0.0 | 0.0 | 0.0 |

### Slide 2
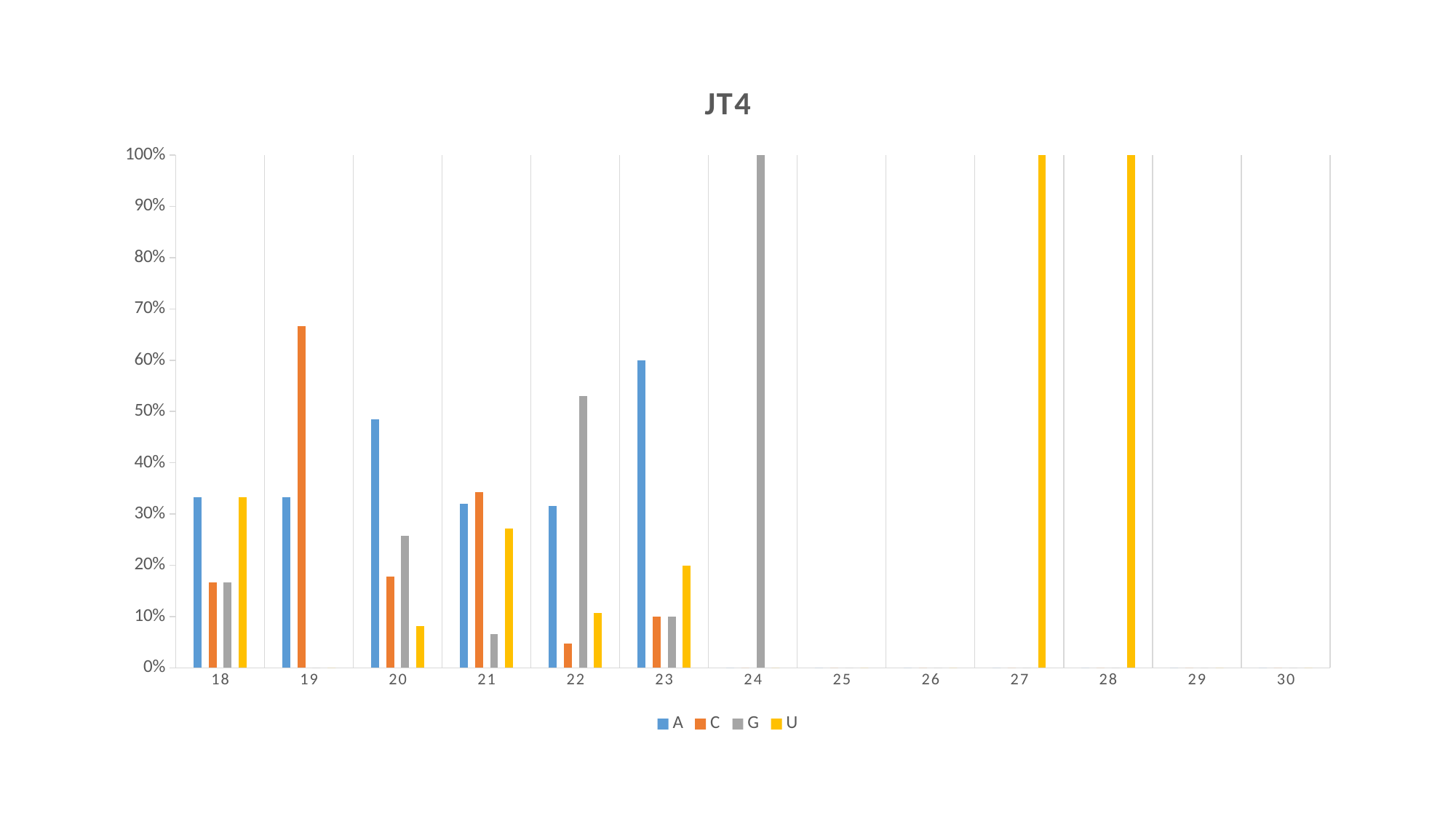

#### Chart: JT4
| Category | A | C | G | U |
|---|---|---|---|---|
| 18 | 0.3333333333333333 | 0.16666666666666666 | 0.16666666666666666 | 0.3333333333333333 |
| 19 | 0.333335 | 0.6666650000000001 | 0.0 | 0.0 |
| 20 | 0.48412499999999997 | 0.17777833333333334 | 0.25714333333333333 | 0.08095333333333334 |
| 21 | 0.3204888888888889 | 0.3420911111111111 | 0.06540333333333333 | 0.27201777777777775 |
| 22 | 0.3156566666666667 | 0.04761833333333334 | 0.5299433333333333 | 0.10678333333333333 |
| 23 | 0.6 | 0.1 | 0.1 | 0.2 |
| 24 | 0.0 | 0.0 | 1.0 | 0.0 |
| 25 | 0.0 | 0.0 | 0.0 | 0.0 |
| 26 | 0.0 | 0.0 | 0.0 | 0.0 |
| 27 | 0.0 | 0.0 | 0.0 | 1.0 |
| 28 | 0.0 | 0.0 | 0.0 | 1.0 |
| 29 | 0.0 | 0.0 | 0.0 | 0.0 |
| 30 | 0.0 | 0.0 | 0.0 | 0.0 |

### Slide 3
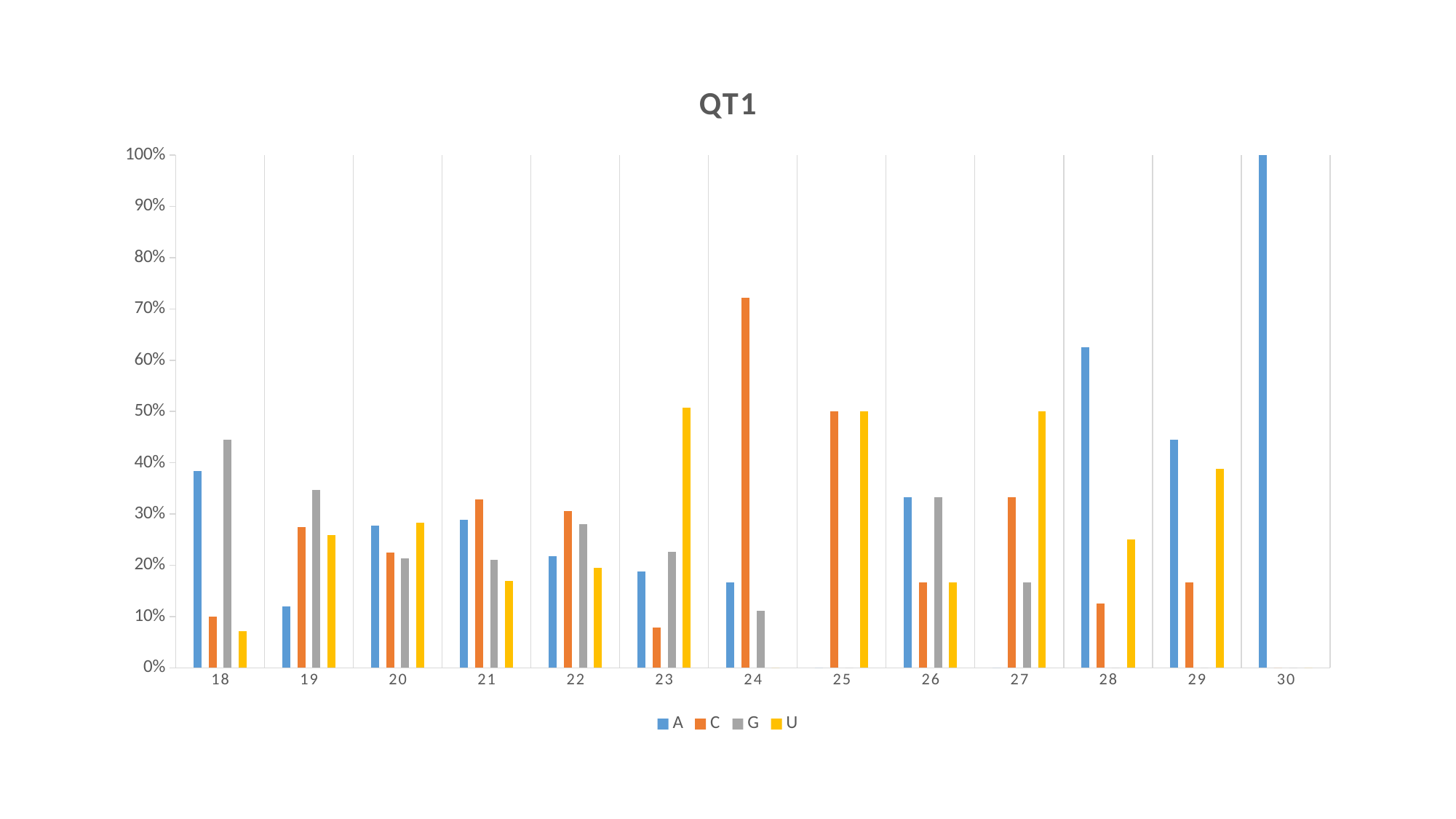

#### Chart: QT1
| Category | A | C | G | U |
|---|---|---|---|---|
| 18 | 0.3833333333333333 | 0.09999999999999999 | 0.4444444444444444 | 0.07222222222222223 |
| 19 | 0.11944444444444442 | 0.27499999999999997 | 0.34722222222222227 | 0.2583333333333333 |
| 20 | 0.2778655555555556 | 0.2251911111111111 | 0.21363999999999997 | 0.2833033333333333 |
| 21 | 0.2895177777777778 | 0.32925999999999994 | 0.21136999999999997 | 0.16985333333333333 |
| 22 | 0.21829444444444443 | 0.30602333333333337 | 0.2803733333333333 | 0.19530777777777777 |
| 23 | 0.18756666666666666 | 0.07830666666666666 | 0.22645555555555555 | 0.5076711111111111 |
| 24 | 0.16666666666666666 | 0.7222222222222222 | 0.1111111111111111 | 0.0 |
| 25 | 0.0 | 0.5 | 0.0 | 0.5 |
| 26 | 0.3333333333333333 | 0.16666666666666666 | 0.3333333333333333 | 0.16666666666666666 |
| 27 | 0.0 | 0.3333333333333333 | 0.16666666666666666 | 0.5 |
| 28 | 0.625 | 0.125 | 0.0 | 0.25 |
| 29 | 0.444445 | 0.16666666666666666 | 0.0 | 0.38888833333333334 |
| 30 | 1.0 | 0.0 | 0.0 | 0.0 |

### Slide 4
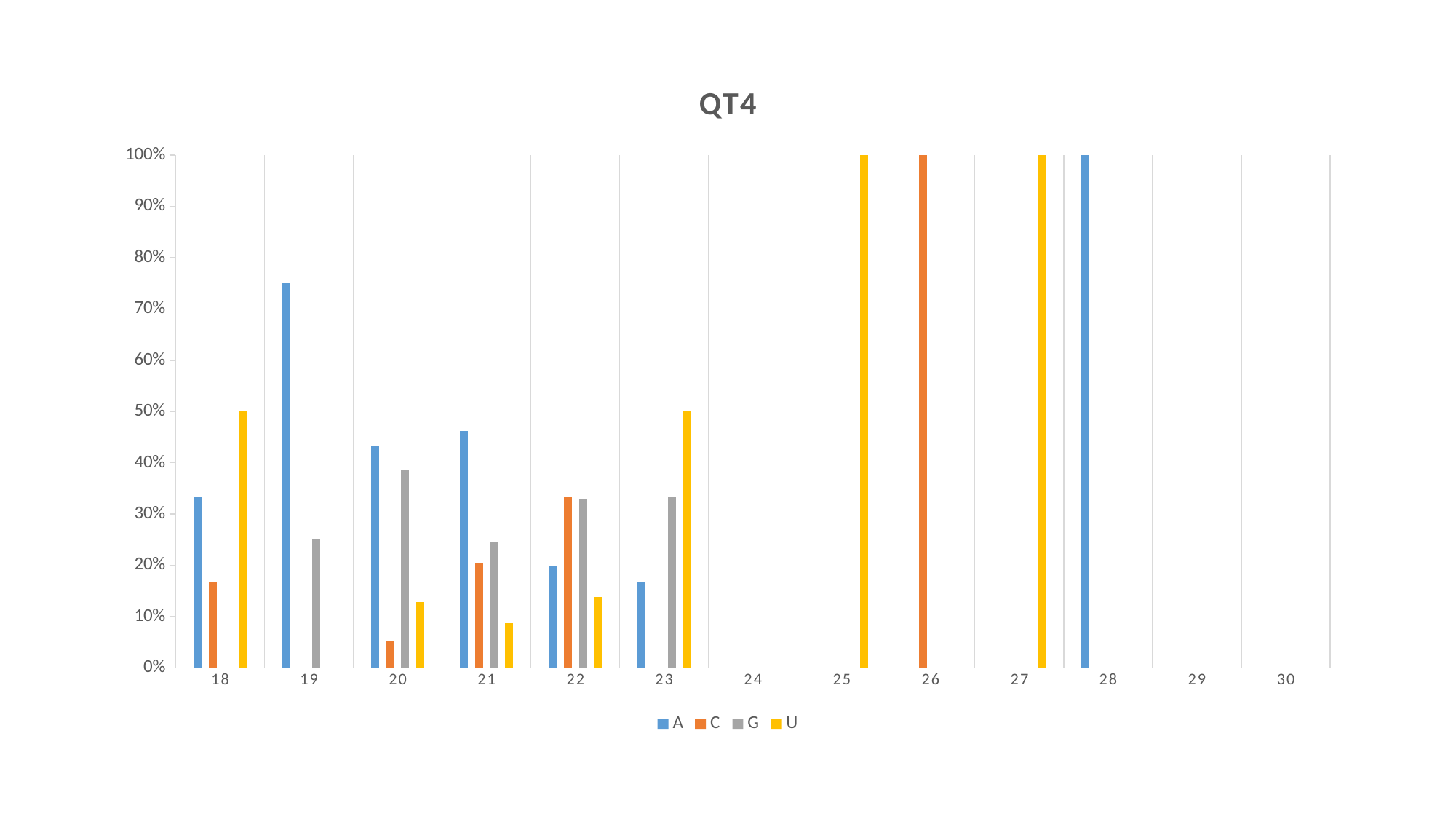

#### Chart: QT4
| Category | A | C | G | U |
|---|---|---|---|---|
| 18 | 0.3333333333333333 | 0.16666666666666666 | 0.0 | 0.5 |
| 19 | 0.75 | 0.0 | 0.25 | 0.0 |
| 20 | 0.4330483333333334 | 0.05128222222222222 | 0.38746444444444444 | 0.1282061111111111 |
| 21 | 0.4621216666666667 | 0.2053016666666667 | 0.24507500000000002 | 0.08750000000000001 |
| 22 | 0.1991111111111111 | 0.3328511111111111 | 0.3296177777777778 | 0.1384188888888889 |
| 23 | 0.166665 | 0.0 | 0.333335 | 0.5 |
| 24 | 0.0 | 0.0 | 0.0 | 0.0 |
| 25 | 0.0 | 0.0 | 0.0 | 1.0 |
| 26 | 0.0 | 1.0 | 0.0 | 0.0 |
| 27 | 0.0 | 0.0 | 0.0 | 1.0 |
| 28 | 1.0 | 0.0 | 0.0 | 0.0 |
| 29 | 0.0 | 0.0 | 0.0 | 0.0 |
| 30 | 0.0 | 0.0 | 0.0 | 0.0 |

### Slide 5
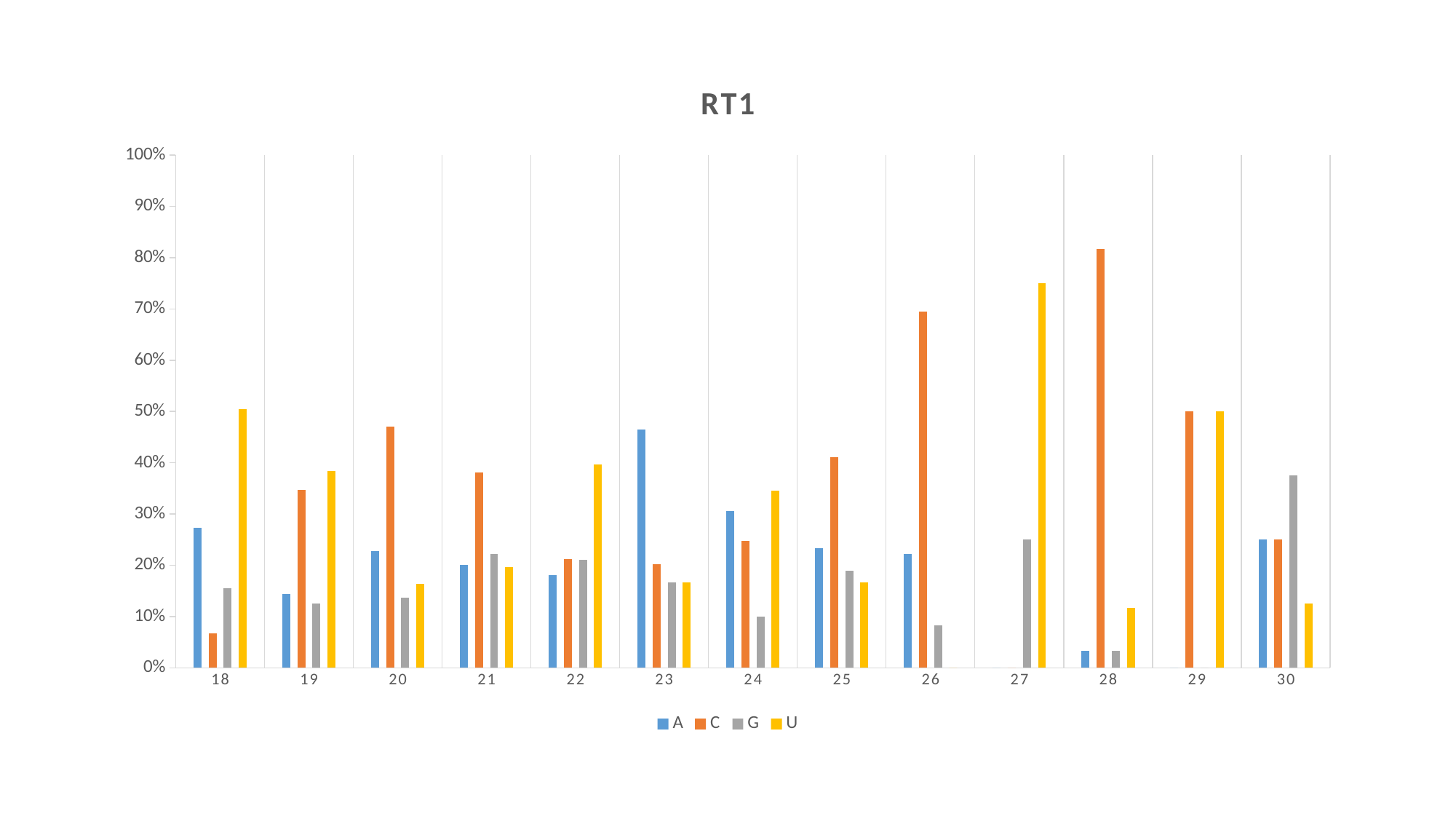

#### Chart: RT1
| Category | A | C | G | U |
|---|---|---|---|---|
| 18 | 0.27381 | 0.06666666666666667 | 0.15476166666666666 | 0.5047616666666667 |
| 19 | 0.1435188888888889 | 0.34722222222222227 | 0.125 | 0.38425888888888887 |
| 20 | 0.22768 | 0.4712077777777777 | 0.13691444444444445 | 0.16419555555555557 |
| 21 | 0.20094444444444448 | 0.38117 | 0.22196888888888888 | 0.19591666666666666 |
| 22 | 0.18130222222222223 | 0.21166666666666667 | 0.2107022222222222 | 0.3963277777777778 |
| 23 | 0.4646722222222222 | 0.20156666666666667 | 0.16709444444444443 | 0.16666666666666666 |
| 24 | 0.3059966666666667 | 0.24779444444444446 | 0.10052888888888889 | 0.34567888888888887 |
| 25 | 0.2333333333333333 | 0.41111111111111115 | 0.18888888888888888 | 0.16666666666666666 |
| 26 | 0.22222166666666668 | 0.694445 | 0.08333333333333333 | 0.0 |
| 27 | 0.0 | 0.0 | 0.25 | 0.75 |
| 28 | 0.03333333333333333 | 0.8166666666666668 | 0.03333333333333333 | 0.11666666666666665 |
| 29 | 0.0 | 0.5 | 0.0 | 0.5 |
| 30 | 0.25 | 0.25 | 0.375 | 0.125 |

### Slide 6
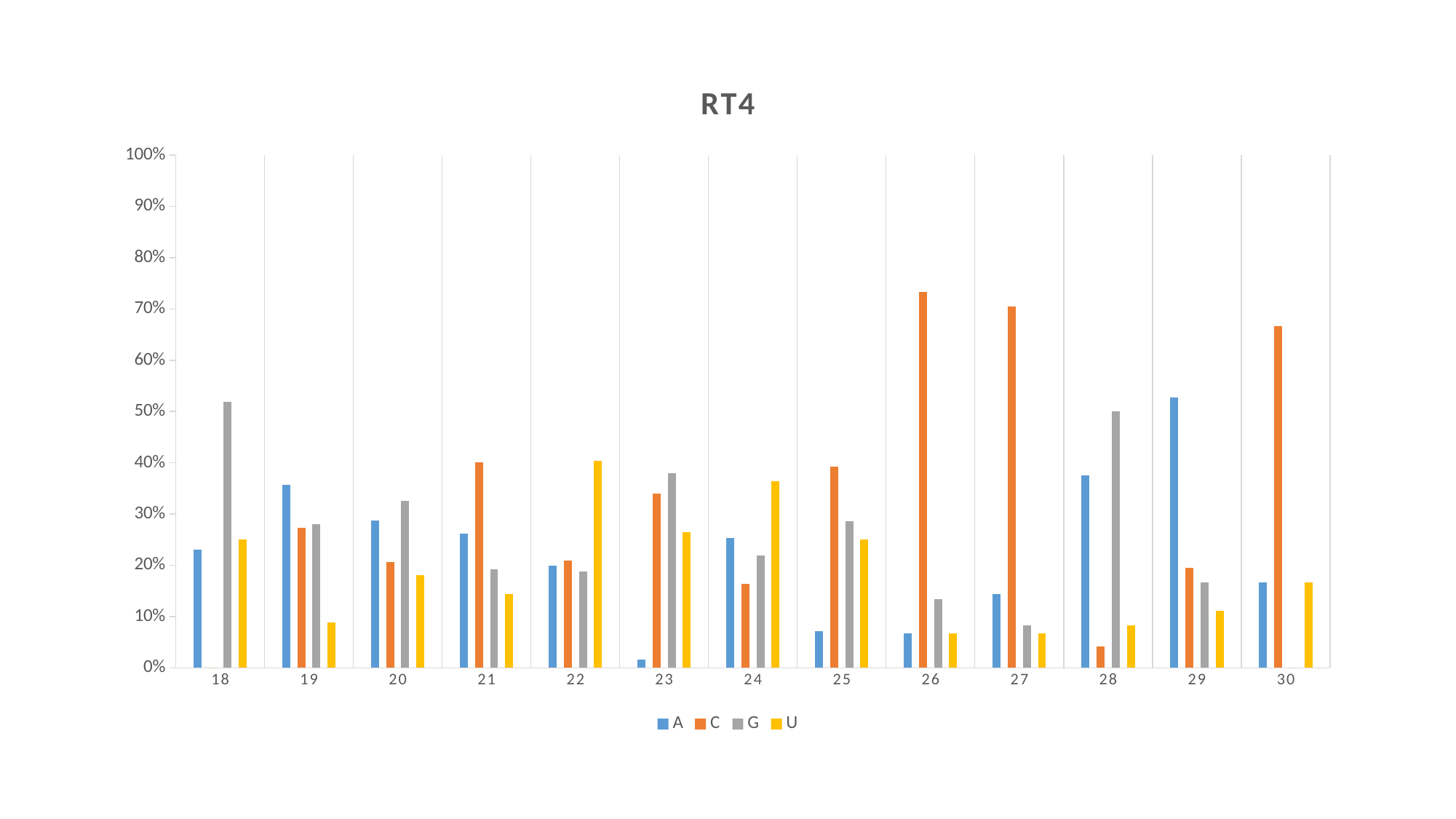

#### Chart: RT4
| Category | A | C | G | U |
|---|---|---|---|---|
| 18 | 0.23077 | 0.0 | 0.51923 | 0.25 |
| 19 | 0.3571416666666667 | 0.2738083333333334 | 0.2797633333333333 | 0.089285 |
| 20 | 0.28807444444444447 | 0.20622777777777776 | 0.3251122222222222 | 0.18058666666666667 |
| 21 | 0.26219 | 0.4004744444444444 | 0.1928122222222222 | 0.14452333333333334 |
| 22 | 0.19912944444444444 | 0.20864777777777777 | 0.1880388888888889 | 0.4041838888888889 |
| 23 | 0.015555 | 0.34 | 0.38000000000000006 | 0.264445 |
| 24 | 0.253245 | 0.16450499999999998 | 0.218615 | 0.36363666666666666 |
| 25 | 0.07143 | 0.392855 | 0.285715 | 0.25 |
| 26 | 0.06666666666666667 | 0.7333333333333334 | 0.13333333333333333 | 0.06666666666666667 |
| 27 | 0.14444333333333334 | 0.7055566666666667 | 0.08333333333333333 | 0.06666666666666667 |
| 28 | 0.375 | 0.0416675 | 0.5 | 0.0833325 |
| 29 | 0.5277766666666667 | 0.19444333333333333 | 0.16666666666666666 | 0.11111 |
| 30 | 0.16666666666666666 | 0.6666666666666666 | 0.0 | 0.16666666666666666 |
