## Supplementary figures and images for "Integration of Transcriptome and Small RNA Sequencing to Decipher Molecular Interaction of *Chenopodium quinoa* Varieties with Cucumber Mosaic Virus"

### Supplementary Figure S1 - PCA plot of DEGs over treatment, time and cultivars.jpg

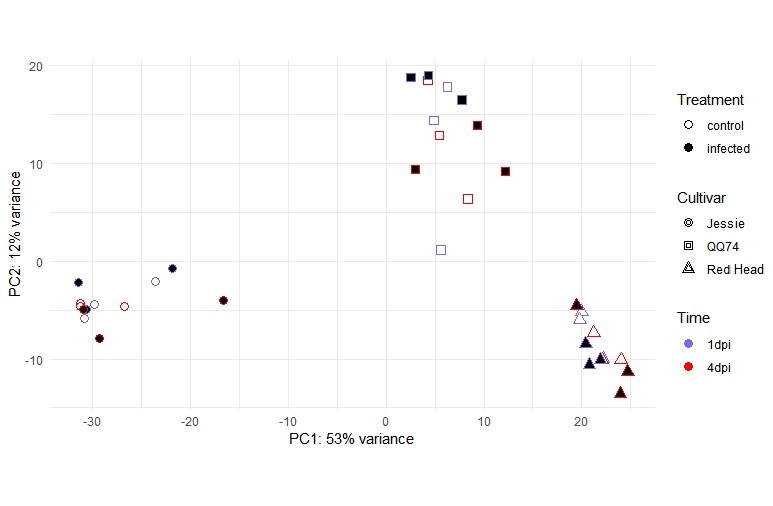

### Supplementary Figure S2 - TSARL1 expression pattern by qPCR.pptx

#### Slide 1
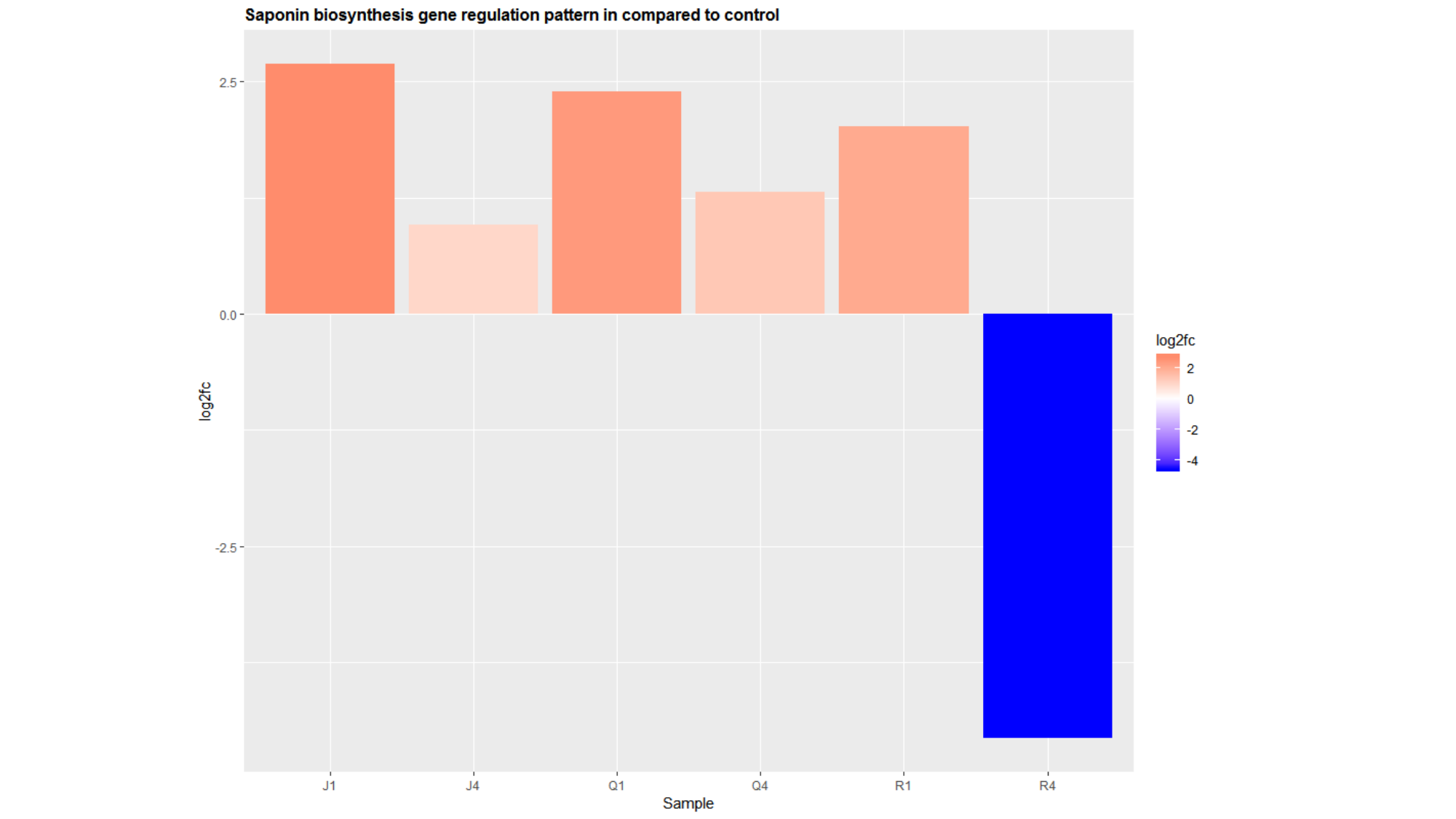
